# High-fidelity 3D live-cell nanoscopy through data-driven enhanced super-resolution radial fluctuation

**DOI:** 10.1101/2022.04.07.487490

**Authors:** Romain F. Laine, Hannah S. Heil, Simao Coelho, Jonathon Nixon-Abell, Angélique Jimenez, Tommaso Galgani, Aki Stubb, Gautier Follain, Samantha Webster, Jesse Goyette, Siân Culley, Guillaume Jacquemet, Bassam Hajj, Christophe Leterrier, Ricardo Henriques

## Abstract

In recent years, the development of new image analysis approaches has highlighted the possibility of recovering superresolution information from short sequences of wide-field images. Our recently developed method, SRRF (Super-Resolution Radial Fluctuations), enables long-term live-cell imaging beyond the resolution limit without specialized hardware. Here, we present eSRRF (enhanced-SRRF), a significant improvement over our initial method, enhancing image fidelity to the underlying structure and resolution. Especially, eSRRF uses automated data-driven parameter optimization, including an estimation of the number of frames necessary for optimal reconstruction. We demonstrate the improved fidelity of the images reconstructed with eSRRF and highlight its versatility and ease of use over a wide range of microscopy techniques and biological systems. We also extend eSRRF to 3D super-resolution microscopy by combining it with multi-focus microscopy (MFM), obtaining volumetric super-resolution imaging of live cells with acquisition speed of ~1 volume/second.

## Introduction

Over the last two decades, super-resolution microscopy (SRM) developments have enabled the unprecedented observation of nanoscale structures in biological systems by light microscopy (1). Stimulated emission depletion (STED) microscopy (2) has led to fast SRM on small fields-of-view with resolution down to 40-50 nm. In contrast, structured illumination microscopy (SIM) (3) provides a doubling in resolution compared to wide-field imaging (~120 nm) with relatively high speed and large fields-of-view. Both superresolution methods rely on complex optical systems to create specific illumination patterns. Single-molecule localization microscopy (SMLM) methods such as (direct) stochastic optical reconstruction microscopy ((d)STORM) (4, 5), photoactivated localization microscopy (PALM) (6) or PAINT (7, 8), take a different approach, exploiting the stochastic ON/OFF switching capabilities of certain fluorescence labeling systems. By separating single emitters in time and sequentially localizing their fluorescence signals, a near molecular resolution (~10-20 nm) can be achieved. However, this commonly requires long acquisition times that range from minutes to days. Image processing and reconstruction tools including multi-emitter fitting localization algorithms (9, 10), Haar wavelet kernel (HAWK) analysis (11), or deep learning assisted tools (12–14) reduce acquisition times by allowing for higher emitter density conditions. Alternatively, fluctuation-based approaches such as super-resolution radial fluctuations (SRRF) (15), super-resolution optical fluctuation imaging (SOFI) (16), multiple signal classification algorithm (MUSICAL) (17) or super-resolution with autocorrelation two-step deconvolution (SACD) (18) can extract super-resolution information from diffraction-limited data (see also Supplementary Table S1). These fluctuation-based approaches only require subtle frame-to-frame intensity variations, rather than the discrete blinking events needed in SMLM, and as such do not require high illumination power densities. Thus, so long as images are acquired with sufficiently high sampling rate to capture spatial and temporal intensity variations, these methods are compatible with most research-grade fluorescence microscopes. This makes them ideally suited for long-term live-cell SRM imaging.

In particular, SRRF is a versatile approach that achieves livecell SRM on a wide range of available microscopy platforms with commonly used fluorescent protein tags (19). It is now a widely used high-density reconstruction algorithm, as highlighted by an important uptake by the community (20–23). The great reception of the SRRF processing tool can also be attributed to its user-friendly implementation, and high accessibility within the Fiji framework (24). However, obtaining optimal reconstruction results with fluctuation-based analysis tools is challenging as they can suffer from reconstruction artifacts and lack signal linearity. Previously, we have developed an approach for the detection and quantification of image artifacts termed SQUIRREL (25). This tool has rapidly become a gold standard in the quantification of super-resolution image quality (26), providing an important platform to assist in the creation of new algorithms, such as those implementing Deep Learning-based methods (27, 28). Here, we present a novel implementation of the SRRF approach termed enhanced SRRF (eSRRF) and highlight its improved capabilities in terms of image fidelity, resolution, and user-friendliness (Figure 1). In eSRRF, we redefined some of the fundamental principles used to estimate radiality and temporal analysis to achieve an improved image quality of the reconstructions. Our new implementation integrates the SQUIRREL engine to provide an automated exploration of the parameter space which identifies the optimal reconstruction parameter set based on quantitative measures of image fidelity and resolution. This optimization is directly driven by the data itself and outlines the trade-offs between resolution and fidelity to the user. By highlighting the optimal parameter range and acquisition configurations, eSRRF minimizes artifacts and non-linearity. Therefore, eSRRF improves overall image fidelity with respect to the underlying structure. The enhanced performance is verified over a wide range of emitter densities and imaging modalities, whilst reducing user bias. We have additionally implemented the capability to achieve true 3D resolution improvement, bypassing the constraints of 2D only of the original SRRF. Obtaining 3D SRM in livecell microscopy still remains a challenge for the field: current implementations of 3D super-resolution methods come at the expense of a limited axial range and long acquisition times, often including major technical demands (29–33). Live-cell super-resolution multidimensional imaging also requires a significantly higher illumination dose than 2D superresolution images, severely compromising cell health and viability (34). In order to implement fluctuation-based live-cell SRM, it is necessary to acquire multiple planes in the axial direction (nearly) simultaneously in order to capture the temporal fluctuations of the emitters in 3D. This has been demonstrated using image splitters (35) but at the cost of additional spherical aberrations. One alternative and powerful approach is multi-focus microscopy (MFM) (36–38), which allows the acquisition of up to 25 planes simultaneously while keeping diffraction-limited image quality in every single plane(39). Here again, as the focal planes are temporally coherent - meaning there is no time lag between axial planes - MFM is ideally well suited for fluctuation-based volumetric live-cell super-resolution imaging, by performing eSRRF on 3D voxels. The estimation of radial fluctuations in 3D using MFM data is a conceptually trivial approach based on the reconstruction principles of SRRF, which however requires a significantly higher computational needs. Here, we demonstrate a full implementation within the Fiji plugin of its capabilities.

**Fig. 1.**
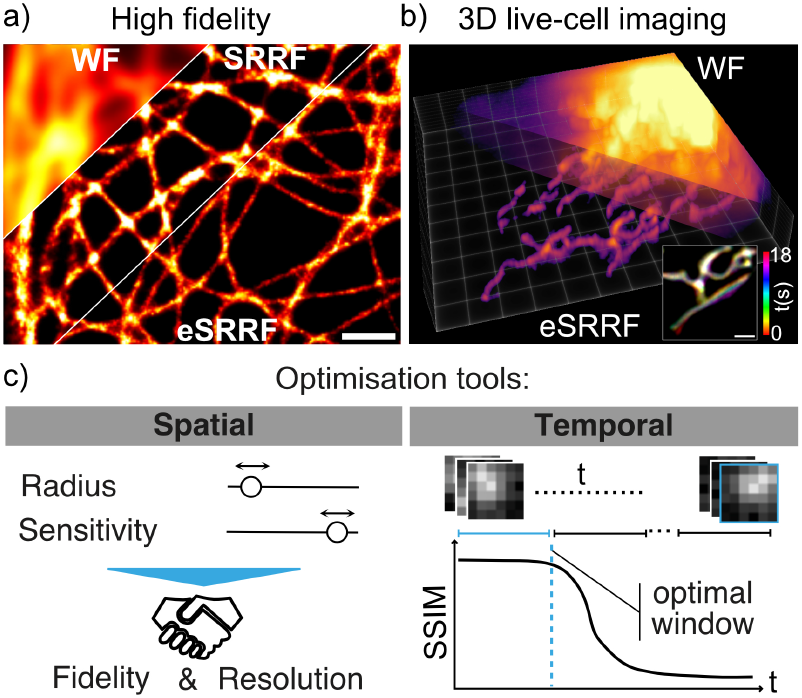
eSRRF achieves high fidelity and 3D live-cell super-resolution. a) The improved reconstruction algorithm of eSRRF surpasses the performance of the original SRRF processing in both image fidelity and resolution (FRC: SRRF 119±32 nm, eSRRF 84±9 nm). b) eSRRF processing also extends to the axial dimension, thus, achieving volumetric live-cell super-resolution imaging in all three dimensions. c) The integrated optimisation toolbox allows to determine optimal reconstruction parameters to maintain high fidelity and resolution (spatial optimisation tool); and to estimate the temporal window size (temporal optimisation tool). Scale bars in a) and the inset in b) 1 μm.

## Results

### eSRRF provides high-fidelity SRM images

Fluctuationbased SRM methods all suffer from the presence of artifacts and/or non-linearity. Here, we designed eSRRF with an emphasis on limiting reconstruction artifacts and maximizing image quality in the reconstruction of super-resolution images. The increased image fidelity results from the implementation of several new and optimized routines in the radial fluctuation analysis algorithm, introduced through a full rewriting of the code. In eSRRF, a raw image time-series with fluctuating fluorescence signals is analyzed (Figure 2a). First, each single frame is upsampled by interpolation (Figure 2b). Here, in contrast to standard SRRF, we introduced a new interpolation strategy. eSRRF now exploits a full data interpolation step based on Fourier transform prior to the gradient calculation. This approach outperforms the cubic spline interpolation employed in the original SRRF analysis by minimizing macro-pixel artifacts (Supplementary Figure S1). Second, following the Fourier transform interpolation, intensity gradients Gx and Gy are calculated and the corresponding weighting factor W based on the user-defined radius R is generated for each pixel. Based on gradient and weighting maps and the user defined sensitivity (S) parameter (Supplementary Table S2), the radial gradient convergence (RGC) is estimated. Thus, in the case of eSRRF, this estimation is not just based on a set number of points at a specific radial distance as it was handled by the previous implementation of SRRF, but over the relevant area around the emitter. This area and how each point contributes to the RGC metric is defined by the W-map. This allows to cover the size of the Point Spread Function (PSF) of the imaging system and, thus, to exploit the local environment of the pixel of interest much more efficiently. Auto- and/or cross-correlation of the resulting RGC time series allows reconstructing a superresolved image which shows high fidelity with respect to the underlying structure (Figure 2b). Compared to the original SRRF, our new eSRRF approach demonstrates a clear improvement in image quality (Figure 2c, Supplementary Figure S2 & S3 and Supplementary Note 1). Although these new implementations make eSRRF processing computationally more demanding, the implementation of OpenCL to parallelise calculations and minimize processing time allows the use of all available computing resources regardless of the platform used.

**Fig. 2.**
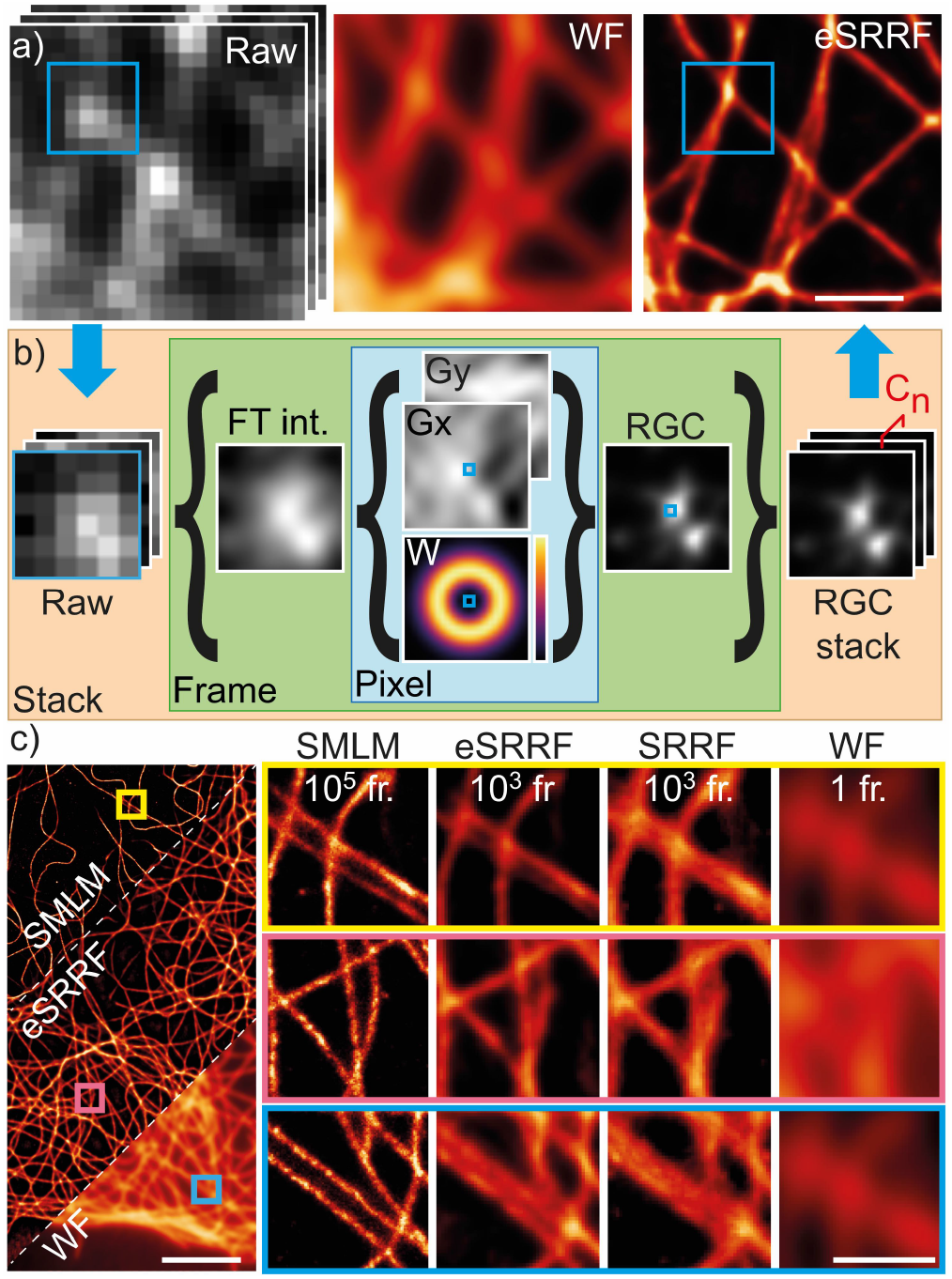
eSRRF image reconstruction produces high-fidelity images. a) eSRRF processing based on a raw data image stack (Raw, left) of a microtubule network allows to surpass the diffraction limited wide-field (WF, middle) image resolution and to super-resolve features that were hidden before (eSRRF, right). b) eSRRF reconstruction steps: Each frame in the stack is interpolated (Fourier transform interpolation, FT int.), from which the gradients *Gx* and *Gy* are calculated. The corresponding weighting factor map *W* is created based on the set radius R. Based on this, the radial gradient convergence (RGC) is calculated for each pixel to compute the RGC map. The RGC stack is then compressed into a super-resolution image by cross-correlation (*C_n_*). c) Super-resolved reconstruction images from eSRRF and SRRF obtained from 1000 frames of high density fluctuation data, created in-silico from an experimental sparse emitter dataset (DNA-PAINT microscopy of immunolabeled microtubules in fixed COS7 cells). The SMLM reconstruction obtained from the sparse data and the wide-field equivalent are shown for comparison.The number of frames used for reconstruction is indicated in each column (FRC resolution estimate: SMLM 71 ± 2 nm, eSRRF 84±11 nm, SRRF 112±40 nm, WF 215 ± 20 nm).

To evaluate the fidelity of eSRRF with respect to the underlying structure, we performed analysis of a DNA-PAINT dataset with sparse localizations. For DNA-PAINT, standard SMLM localization algorithms applied to the raw, sparse data can provide an accurate representation of the underlying structure. By temporally binning the raw data, we generated a high-density dataset comparable to a typical live-cell imaging acquisition. Figure 2c shows the comparison of the ground-truth (SMLM), eSRRF, SRRF and equivalent wide-field (WF) data. eSRRF is in good structural agreement with the ground truth and shows a clear resolution improvement over both the wide-field and the SRRF reconstruction. Line profiles reveal that eSRRF resolves features that were only visible in the SMLM reconstruction (Supplementary Figure S2). This observation is supported by the estimation of the image resolution by Fourier Ring Correlation (FRC) (41), which provides a quantitative assessment of the performance of the different image reconstruction modalities. Furthermore, the enhanced image fidelity recovered from eSRRF is quantitatively confirmed using SQUIRREL analysis on both simulated and experimental data (Supplementary Figures S3 & S4, Supplementary Note 1).

eSRRF not only achieves higher fidelity in image reconstruction than SRRF, but also provides a robust and reproducible reconstruction method over a wide range of emitter densities. To estimate the range of emitter densities compatible with eSRRF, we again use low-density DNA-PAINT acquisitions and temporally binned the images with varying numbers of frames per bin. By increasing the number of frames per bin, the density of molecules in each binned frame increases. Whilst the total number of molecules remains consistent throughout, this approach allows us to monitor the performance of eSRRF as a function of emitter density. Supplementary Figure S5 presents the results from this analysis across the 3 temporal analyses provided (AVG, TAC2, VAR, see Supplementary Note 2). For each density range, a specific set of the processing parameters will provide the best image quality (see Supplementary Table S2 & Supplementary Notes 2) allowing access to high fidelity super-resolved image reconstructions across a wide range of experimental conditions. At high emitter densities, eSRRF also outperforms the high-density emitter localization algorithm of ThunderSTORM (9) even in combination with HAWK analysis (11) (Supplementary Figure S6). At low densities, single emitter fitting still provides unsurpassed localization precision and image resolution; however, it requires the processing of a large number of images. Here, eSRRF can provide a fast preview of the reconstructed image (Supplementary Figure S7).

### eSRRF provides an adaptable reconstruction parameter exploration scheme

The decision to use a specific set of parameters for an image reconstruction is often based on user bias. This can lead to the inclusion of artifacts in the data (11). In order to alleviate user bias and artefacts, here, we develop a quantitative reconstruction parameter search based on SQUIRREL. For this we compute visual maps of the FRC resolution and image fidelity as a function of radius *R* and sensitivity *S*, exploring the eSRRF reconstruction parameter space. We use FRC to determine image resolution and Resolution-scaled Pearson correlation coefficient (RSP) as a metric for image fidelity. The approach highlights the trade-offs between FRC resolution and RSP fidelity (Supplementary Movie M1) done as a consequence of reconstruction parameter choice. In order to balance the two metrics, we use an F1-calculation to compute the QnR score:

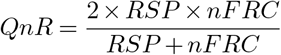

Here *nFRC* is the normalized FRC resolution metric, ranging between 0 and 1, with 0 representing a poor resolution and 1 representing a high resolution.

The QnR score ranges between 0 and 1, where scores close to 1 represent a good combination of FRC resolution and RSP fidelity, whereas a QnR score close to 0 represents a low-quality image reconstruction. Figure 3 shows a representative dataset acquired with COS7 cells expressing lyn kinase – SkylanS previously published by Moeyaerd et al. (40, 42). The eSRRF parameter scan analysis (Figure 3a) shows how the RSP fidelity and the FRC resolution are affected by reconstruction parameters. RSP fidelity is high when using a low sensitivity and/or low radius. In contrast, FRC resolution improves upon increasing the sensitivity over a large range of radii. This can be explained by the appearance of nonlinear artifacts at high sensitivity leading to low RSP fidelity but high FRC resolution. In addition, as the radius increases, the resolution of the reconstructed image decreases. The QnR metric map, shown in Figure 3b, demonstrates that a balance can be found that leads to both a good resolution and a good fidelity. Figure 3c shows a range of image reconstruction parameters: the optimal reconstruction parameter set (R=1.5, S=4, Figure 3c, i) and two other suboptimal parameter sets (Figure 3c ii and iii). Figure 3c ii shows a low-resolution image, whereas Figure 3c iii has a high level of non-linearity - the result of an inappropriately high sensitivity. Our automated parameter search implemented in eSRRF enables the user to find the optimal settings for the specific dataset being analyzed. This makes eSRRF not only user-friendly but also ensures reproducible results with minimized user bias.

**Fig. 3.**
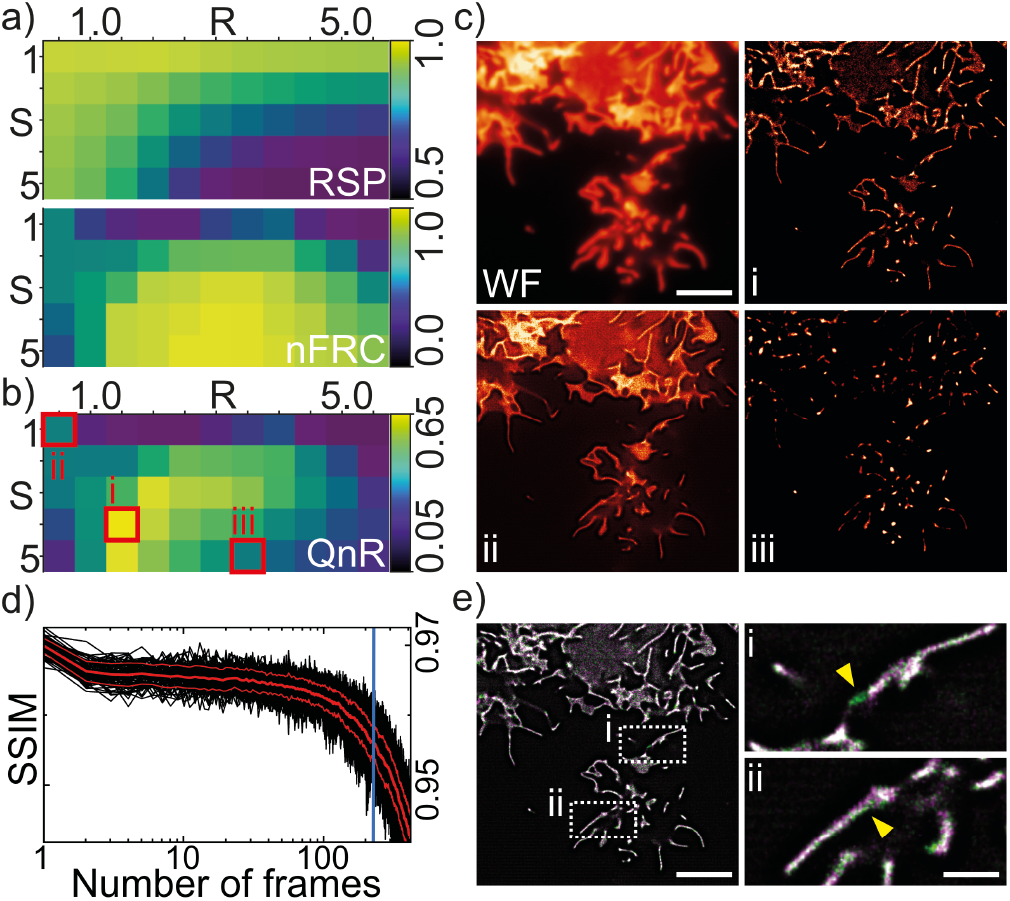
eSRRF provides an automated reconstruction parameter search. a-c) Finding the optimal parameters to calculate the RGC. a) RSP and FRC resolution maps as functions of Radius (R) and Sensitivity (S) reconstruction parameters for a live-cell TIRF imaging dataset published by Moeyaert et al. (40). The COS-7 cells are expressing the membrane targeting domain of Lyn kinase – SkylanS and were imaged at 33 Hz. b) Combined QnR metric map showing the compromise between fidelity and FRC resolution. c) Wide-field image, optimal eSRRF reconstruction (i, *R*=1.5, S=4), low resolution reconstruction (ii, *R*=0.5, S=1) and low fidelity reconstruction (iii, R=3.5, S=5). d-e) Estimating the optimal time window for the eSRRF temporal analysis based on tSSIM. d) The SSIM metric is observed over time, after ~200 frames it displays a sharp drop. The optimal time window is marked by the blue line. e) A color-overlay of two consecutive reconstructed eSRRF frames with the optimal parameters and a frame window of 200 frames displays significant differences between the structures (marked in i and ii), which would lead to motion blurring in case of a longer frame window. Scale bars in c) and e) 20 μm, in the e-i) and ii) 5 μm.

An important aspect of live-cell super-resolution imaging is its capacity for observation and quantification of dynamic processes at the molecular level. To address this, we have further integrated temporal structural similarity (tSSIM) analysis. Here, we calculate the progression of the structural similarity (43) at the different time points of the image stack relative to the first frame (Supplementary Figure S8). This allows us to identify the local molecular dynamics (Supplementary Figure S9) and estimate the maximum number of frames within which the structural similarity is retained - meaning there is no observable movement (Figure 3 d)). By combining tSSIM with eSRRF, we can determine the optimal temporal sampling rate required to recover such dynamics, whilst avoiding motion blur artifacts (Figure 3 e).

### eSRRF works across a wide range of live-cell imaging modalities

Here, we test our approach on a wide range of imaging modalities including total internal reflection fluorescence (TIRF), fast highly inclined and laminated optical sheet (HiLO)-TIRF (44), spinning-disk confocal (SDC) (45) and lattice light-sheet (LLS) microscopy (46). We show that eSRRF provides high quality live-cell SRM images (Figure 4). First, we imaged ffDronpa-MAP4 in live HeLa cells using TIRF microscopy. eSRRF reconstruction allows for a superresolved view of the microtubule network in living cells (Figure 4a). Second, we evaluated the dynamic rearrangement of the endoplasmic reticulum (ER) in living COS-7 cells. Acquired using HiLO-TIRF, the fast acquisition rates allowed us to track the ER network tagged with PrSS-mEmerald-KDEL, at super-resolution level (Figure 4b). Here, a sampling rate of 50 ms is achieved using rolling window analysis of eSRRF (Supplementary Figure S10 & Supplementary Movie M2). While TIRF and HiLO-TIRF imaging are set out for fast high contrast imaging in close vicinity to the coverslip surface, SDC excels in fast and gentle in vivo imaging. This entails imaging far away from the coverslip and deep inside challenging samples as spheroids and live organisms where eSRRF achieves enhanced performance as well (Supplementary Figure S11).

**Fig. 4.**
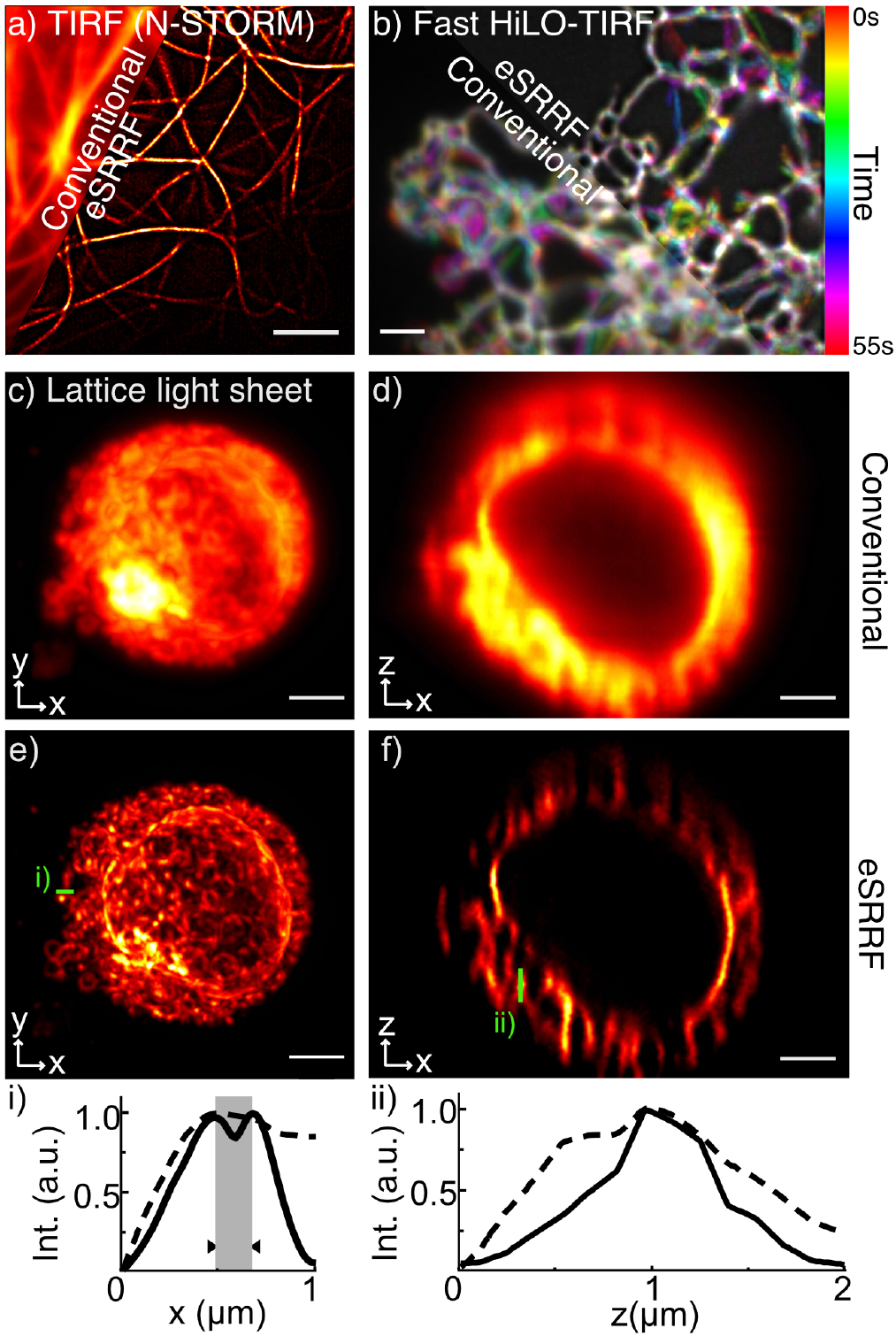
Applications of eSRRF to a range of imaging modalities. a) Live-cell TIRF imaging of HeLa cells expressing ffDronpa-MAP4 (FRC resolution Conventional/eSRRF: 266±51/128±51 nm). b) Live-cell HiLO-TIRF of COS-7 cells expressing PrSS-mEmerald-KDEL marking the ER lumen. The temporal evolution is color-coded (FRC resolution Conventional/eSRRF: 254±11/143±56 nm). c-f) Lattice light sheet (LLS) imaging of the ER in live Jurkat cells at a rate of 7.6 mHz per volume (79 x 55 x 35 μm3). c, xy projection, and d, xz projection using LLS microscopy. e, xy projection, and f, xz projection using eSRRF reconstruction. As the acquisition, the eSRRF processing was applied on a slice-by-slice basis. Line profiles corresponding to the x- and z-direction are shown in i and ii, respectively. The line profile in i reveals sub-diffraction features separated by 190 nm (marked in gray) (FRC resolution LLS/eSRRF: 164±9/84±43 nm). Scale bar in a) 5 μm, b) 2 μm and c-f) 3 μm.

eSRRF can also be applied to volumetric live-cell datasets as obtained for example with LLS microscopy (Supplementary Movie M3). Here, the eSRRF reconstruction of volumetric image stacks is obtained by processing each slice sequentially. Note that this approach can only effectively improve the lateral resolution (x-y plane), while there is a sharpening comparable to deconvolution in the z-direction, no resolution improvement over the diffraction limited images should be expected. Figure 4 c-f) shows the plane by plane eSRRF processing of a LLS dataset of the ER in live Jurkat cells allows to distinguish sub-diffraction limited features along the x-direction (Figure 4e), line profile i)), but not in the z-direction (Figure 4f), line profile ii)).

### 3D live-cell super-resolution imaging by eSRRF in combination with multifocus microscopy

3D imaging capability is becoming increasingly important to understand molecular dynamics and interactions within the full context of their environment. In particular, obtaining true 3D SRM with improved resolution along the axial direction has recently become a key focus of development in the field. Fluctuation-based SRM approaches have also been extended to 3D, notably SOFI (16, 35) and, more recently, random illumination microscopy (RIM) (47), an approach that combines the concepts of fluctuation microscopy and the SIM demodulation principle. To realize 3D eSRRF, we extended the algorithm to calculate the RGC in 3D. Consequently, we can reconstruct a volumetric image with enhanced resolution in the axial direction and in the lateral image plane. The approach was first validated with a simulated 3D dataset (see Supplementary Figure S12). In practice, extending fluctuationbased analysis methods to 3D requires the near-simultaneous detection of multiple planes. To determine the radial symmetry in 3D, within the time scale of the temporal fluctuations of the fluorescent probes, the whole volume needs to be acquired concurrently. MFM allows for the detection of multiple axial planes onto a single camera at the same time by using an aberration corrected diffractive optical element (36–38) (Supplementary Figure S13). By combining MFM and eSRRF, a super-resolved volumetric view (20 x 20 x 3.6 μm3) of the mitochondrial network architecture and dynamics in U2OS cells was acquired at a rate of 1Hz (Figure 5). The eSRRF processing achieved super-resolution in lateral and axial dimensions, revealing sub-diffraction limited structures. Figure 5 shows that eSRRF reveals structures which would otherwise remain undetected using conventional MFM, even after applying deconvolution (Figure 5 i-ii) & Supplementary Figure S14). By processing the entire image sequence the dynamic fluctuations of the mitochondrial network in the living cell can be observed at super-resolution level over time (Figure 5 iii), Supplementary Movie M4 & M5).

**Fig. 5.**
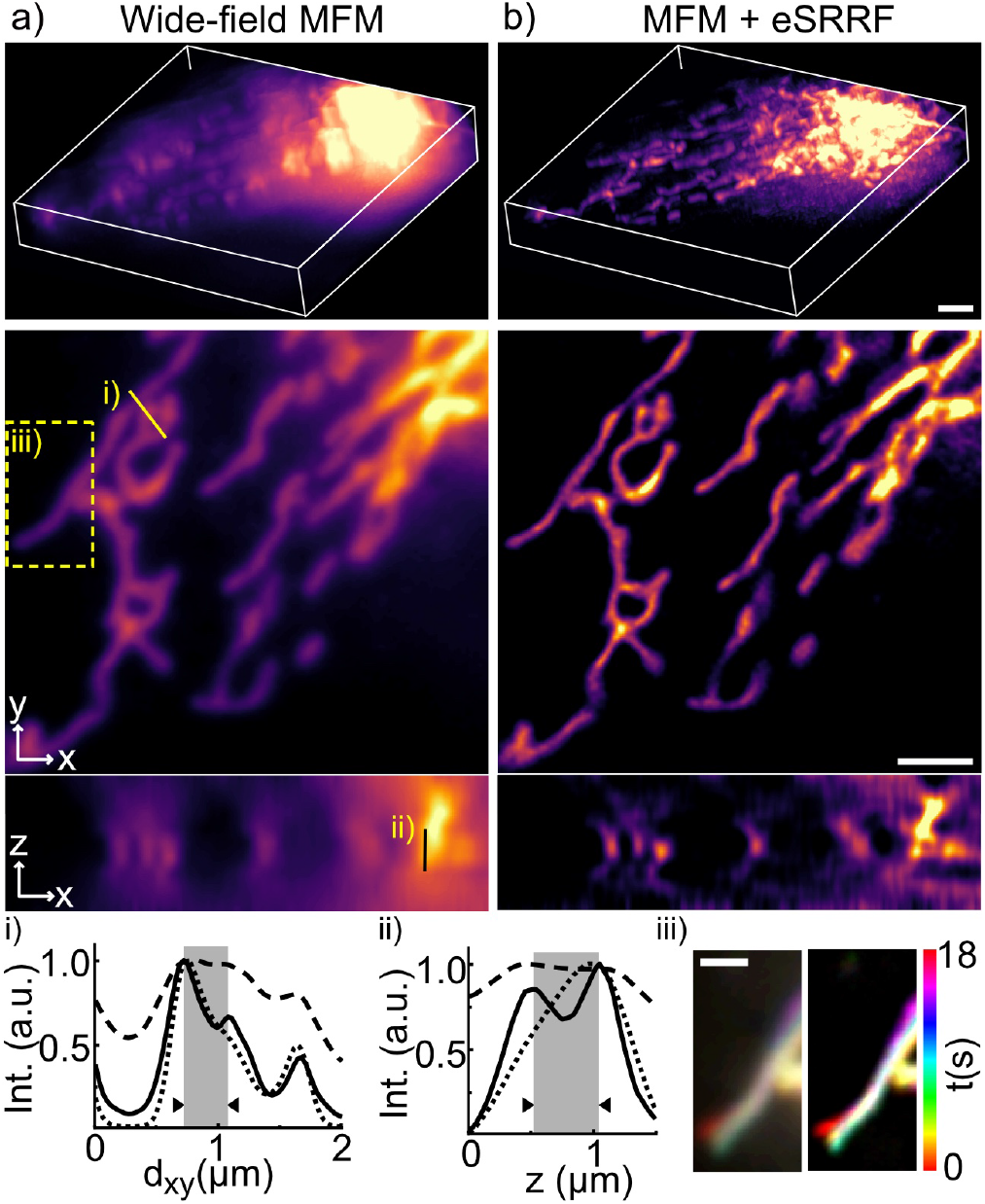
SRRF and MFM allows 3D live-cell super-resolution. a) Live-cell volumetric imaging in MFM widefield configuration of U2OS cells expressing TOM20-Halo, loaded with JF549. b) 3D eSRRF processing of the dataset creates a superresolved volumetric view of 20 x 20 x 3.6 μm3 at a rate of 1Hz (MFM + eSRRF). a-b) Top: 3D rendering, middle: single cropped z-slice (FRC resolution in xy: interpolated: 231 ± 10 nm, eSRRF: 74 ± 12 nm), bottom: single cropped y-slice (xz eSRRF: 173 ± 19 nm) with i) and ii) mark the positions of the respective line profiles in x,y and z-plane in the MFM (dashed line), deconvolved MFM (dotted line, see Supplementary Figure S14) and MFM+eSRRF (solid line) images. The distance of the structures resolved by eSRRF processing (marked gray) is 360 nm in the lateral directions (x,y) and 500 nm in the axial direction (z). iii) marks the displayed area of the temporal color coded projection of a single z slice over the whole MFM (left) and MFM+eSRRF (right) acquisition. Scale bars 2 μm in a-b) and 1 μm in iii).

## Conclusion

The new eSRRF approach builds on the previous capacity of SRRF, considerably improving the method image reconstruction quality and fidelity. It showcases a novel analytical engine for calculating the Radial Gradient Convergence transform, replacing the lower quality Radiality transform of the original SRRF method. These modifications have also allowed us to extend the approach into full 3D superresolution, by combining it with multi-focus microscopy. eSRRF also introduces a data-driven parameter optimization approach that aids users in selecting optimal parameters learned directly from the data to be analyzed. These optimal parameters are chosen by balancing the need for high reconstruction fidelity together with high spatial and temporal resolution. While we demonstrate this original concept in eSRRF, we expect this strategy to be easily transferable to other superresolution methods that require an analytical component, as is the case for SMLM approaches. To demonstrate the broad applicability of eSRRF, we showcase its application to a wide range of biological samples from single cells to organisms, and imaging techniques from widefield, TIRF, light sheet, spinning-disc confocal, and SMLM imaging modalities. eSRRF shows robust performance over the different signal fluctuation dynamics displayed by various organic dyes and fluorescent proteins and over a wide range of marker densities, recovering high fidelity super-resolution images even in challenging conditions in which single-molecule algorithms will fail. To achieve optimal spatial and temporal resolution, minimize reconstruction artifacts and reduce user bias, we have implemented a metric for image resolution and fidelity, which we call QnR, used to perform data-driven parameter optimization alongside temporal window optimization. This makes eSRRF a super-resolution method that learns and guides users on how to best analyze their data, providing essential information to find ideal phototoxicity-sensitive livecell super-resolution imaging conditions. Through these developments, eSRRF provides novel fundamental principles to make live-cell SRM more robust and reliable. eSRRF is implemented as an open-source GPU accelerated Fiji plugin, accompanied by detailed [user guide](https://github.com/HenriquesLab/NanoJ-eSRRF), making it widely available to the bioimaging community.

## Availability

eSRRF is available as Supplemental Software or can be accessed from our [GitHub page](https://github.com/HenriquesLab//NanoJ-eSRRF). This resource is fully open-source and includes a Wiki manual.

## Data availability

Example datasets are available on our [Zenodo (110.5281/zenodo.6466472)](https://doi.org/10.5281/zenodo.6466472). Further data are available on request.

## ACKNOWLEDGEMENTS

We thank Jennifer Lippincott-Schwartz and Christopher Obara, Janelia Farm, for their assistance with the dynamic ER-TIRF data and for reading the manuscript, Peter Dedecker, University of Leuven, for providing the ffDronpa-MAP4 plasmid. This work was supported by the Gulbenkian Foundation (H.S.H., S.C. (Coelho), R.H.) and received funding from the European Research Council (ERC) under the European Union’s Horizon 2020 research and innovation programme (grant agreement no.101001332 to R.H.), the European Molecular Biology Organization (EMBO-2020-IG-4734 to R.H. and ALTF 499-2021 to H.S.H), the Wellcome Trust (203276/Z/16/Zto R.H.), the Fundação para a Ciência e Tecnologia, Portugal, (FCT fellowship CEECIND/01480/2021 to H.S.H., CEECIND/07466/2022 and PTDC/08248/2022 to S.C. (Coelho)), the Chan Zuckerberg Initiative Visual Proteomics Grant (vpi-0000000044 to R.H.), InnOValley Proof of Concept Fund (IOVPoC-2021-01 to S.C. (Coelho)) and National Health and Medical Research Council (NHMRC) of Australia (APP1183588 to S.C. (Coelho) and J.G.). R.F.L. would like to acknowledge the support of the MRC Skills development fellowship (MR/T027924/1). S.C. (Culley) would like to acknowledge support from a Royal Society University Research Fellowship. This study was supported by the Academy of Finland (G.J., 338537 and G.F., 332402), the Sigrid Juselius Foundation (G.J.), the Cancer Society of Finland (G.J.), the Åbo Akademi University Research Foundation (G.J., CoE CellMech), the Drug Discovery and Diagnostics strategic funding to Åbo Akademi University (G.J.). The Cell Imaging and Cytometry Core facility (Turku Bioscience, University of Turku, Åbo Akademi University, and Biocenter Finland) and Turku Bioimaging are acknowledged for services, instrumentation, and expertise. C.L. would like to acknowledge funding of CNRS through the ATIP (A0 2016) grant. B.H. acknowledges funding from the Fondation pour la recherche médicale (FRM; DEI20151234398), the Agence National de la recherche (ANR-19-CE42-0003-01), the LabEx CelTisPhyBio (ANR-11-LABX-0038, ANR-10-IDEX-0001-02), the Institut Curie, Agence pour la Recherche sur le Cancer (ARC Foundation), DIM ELICIT and from ITMO Cancer of Aviesan on funds administered by Inserm (grant N°20CP092-00). B.H. recognizes the support of France-BioImaging infrastructure grant ANR-10-INBS-04 (Investments for the future).

## EXTENDED AUTHOR INFORMATION

- Romain F. Laine: LaineBioImaging
- Hannah S. Heil: Hannah_SuperRes
- Simao Coelho: simaopc
- Jonathon Nixon-Abell: AbellJonny
- Angélique Jimenez
- Tommaso Galgani:
- Aki Stubb: akistub
- Gautier Follain: Follain_Ga
- Samantha Webster:
- Jesse Goyette:
- Siãn Culley: SuperResoluSian
- Guillaume Jacquemet: guijacquemet
- Bassam Hajj: Bassam_A_HAJJ
- Christophe Leterrier: christlet
- Ricardo Henriques: HenriquesLab

## AUTHOR CONTRIBUTIONS

R.F.L., B.H., C.L and R.H. designed the study. R.F.L., S.C. (Culley), and R.H. wrote the algorithm. S.C. (Culley) wrote the script for data simulation. R.F.L., S.C. (Coelho), H.S.H., J.N-A., A.J., T.G., A.S., G.F., G.J., B.H. and C.L. performed the experiments. H.S.H., tested, documented, and validated the algorithm, and analyzed the data. S.C. (Coelho) analyzed the LLS data. All authors planned experiments and contributed to the writing of the article.

## COMPETING FINANCIAL INTERESTS

The authors declare no competing financial interests.

## Online Methods

The imaging conditions and eSRRF processing parameters for each data set are summarized in S3.

### Fluorescence microscopy simulations

The field of simulated fluorescent molecule distribution and the MFM dataset of stacked lines was simulated over a 5 nm resolution grid with the NanoJ-eSRRF>Fluorescence simulator application. Each fluorescent emitter was allowed to blink independently with on/off rate of 100 s^-1^ and 50 s^-^1 respectively over the entire acquisition without bleaching (500 frames at 10 ms exposure). The simulated fluorescence image produced by this distribution of beads was created by convolution with a Gaussian kernel with *σ* = 0.21 *λ/NA*, as suggested by Zhang et al. (1)), where *λ* is the emission wavelength (here 580 nm) and *NA* is the numerical aperture of the microscope (here *NA* = 1.4). A pixel size of 100 nm was chosen for the final fluorescence image, in agreement with our experimental set-up. A realistic Poisson photon noise and a Gaussian read-out noise were added to the images in order to simulate an experimental dataset. The diffusing particle datasets were generated as single emitters represented by a Gaussian PSF and with Gaussian noise moving at constant speed with a Python script available as GoogleCoLabs Jupyter notebook on [GitHub](https://github.com/HenriquesLab//NanoJ-eSRRF).

### DNA-PAINT of microtubule network

COS-7 cells (ATCC CRL-1651) were cultured in phenol free DMEM (Gibco) supplemented with 2 mM GlutaMAX (Gibco), 50 U/ml penicillin, 50 μg/ml streptomycin (Penstrep, Gibco) and 10% foetal bovine serum (FBS; Gibco) at 37 °C humidified incubator with 5% CO2. Cells were seeded on ultraclean (2) 18 mm diameter thickness 1.5 H coverslips (Marienfeld) at a density of 0.3–0.9 × 105 cells/cm2. Cells were fixed and stained according to previously published protocols (3). Cells were extracted at 37 °C for 45 s in 0.25% Triton-X (T8787, Sigma), 0.1% glutaraldehyde in the cytoskeleton-preserving buffer “PIPES-EGTA-Magnesium” (PEM: 80 mM PIPES pH 6.8, 5 mM EGTA, 2 mM MgCl2) followed by 10 min in 0.25% Triton-X, 0.5% glutaraldehyde in PEM. After a 7 min quenching step with a fresh solution of 0.1% NaBH4 in phosphate buffer at room temperature cells were permeabilized and blocked for 1.5h at room temperature in blocking buffer (phosphate buffer 0.1 M pH 7.3, 0.22% gelatin (G9391, Sigma), 0.1% Triton X-100). Primary Antibody labeling was performed at 4 °C overnight with a mix of two anti-α-tubulin mouse monoclonal IgG1 antibodies (DM1A (T6199, Sigma) and B-5–1-2 (T5168, Sigma) diluted 1:300 in blocking buffer. After 3×10 min washes with blocking buffer, the cells are incubated with a goat anti-mouse antibody conjugated to a DNA sequence (P1 docking strand, Ultivue kit) for 1h at room temperature diluted at 1:100 in blocking buffer. After incubation, cells are washed with blocking buffer for 10 min and 2x 10 min with phosphate buffer. DNA-PAINT imaging was performed on a N-STORM microscope (Nikon) equipped with 647 nm lasers (125 mW at the optical fiber output). After injection of 0.25 nM imager strand (I1-ATTO655, Ultivue) in 500 mM NaCl in 0.1 M PBS pH 7.2 buffer, 50,000 frames were acquired at 60 % power of the 647 nm laser with an exposure time of 30 ms/frame to obtain low density ground truth data.

### Live-cell TIRF microscopy of MAP4 in HeLa cells

HeLa cells were plated 8-well Ibidi dishes and transfected with ffDronpa-MAP4 (kind gift of Peter Dedecker) using Lipofectamine 2000 as per manufacturer’s protocol. Cells were imaged in PBS, using the Nikon N-STORM TIRF microscope equipped with 100x TIRF objective and an Andor iXon EM-CCD camera. Cells were continuously imaged at 100 fps (10 ms exposure) in TIRF with a 488 nm laser at an illumination density of 0.65 kW/cm^2^.

### Live-cell HiLO-TIRF microscopy

COS7 cells (ATCC) were grown in phenol red-free Dulbecco’s modified Eagle medium (DMEM) supplemented with 10% (v/v) FBS (Gibco), 2 mM L-glutamine (Thermo), 100 U/ml penicillin and 100 μg/ml streptomycin (Thermo) at 37 °C and 5% CO^2^. 25 mm Number 1.5 coverslips (Warner scientific) were precleaned by: (i) 12-hour sonication in 0.1% Hellmanex™ (Sigma); (ii) five washes in 300ml of distilled water; (iii) 12-hour sonication in distilled water; (iv) an additional round of five washes in distilled water; (v) sterilised in 200 proof ethanol and allowed to air dry. Coverslips were coated with 500μg/ml phenol red free matrigel (Corning). Cells were seeded at 60% confluency. Transfections were performed using Fugene6 (Promega) according to the manufacturer’s instructions. Each coverslip was transfected with 750 ng of PrSS-mEmerald-KDEL to label the ER structure, and with 250 ng of HaloTag-Sec61b-TA (not labelled with ligand for these experiments). Imaging was performed using a customized inverted Nikon Ti-E microscope outfitted with a live imaging chamber to maintain temperature, CO^2^, and relative humidity during imaging (Tokai Hit). The sample was illuminated with a fiber-coupled 488nm laser (Agilent Technologies) through a rear-mount TIRF illuminator. Imaging was performed such that the TIRF angle was manually adjusted below the critical angle to ensure HiLO illuminations and that the ER was captured within the illumination plane. The average power density over the full illumination field was 123 mW/cm2. Fluorescence was collected using a 100x α-Plan-Apochromat 1.49 NA oil objective (Nikon) using a 525/50 filter (Chroma) placed before a iXon3 electron multiplying charged coupled device camera (EM-CCD, DU-897; Andor). Imaging was performed with 5ms exposure times for 60 seconds. The precise timing of each frame was monitored using an oscilloscope directly coupled into the system (mean frame rate 95Hz).

### Sample preparation & acquisition for lattice light sheet microscopy of Jurkat cells

The ER of Jurkat cells were stained with BODIPY ER-Tracker and incubated on a poly-L-lysine-coated 5 mm round coverslip at 37 °C. The lattice light-sheet (LLS) data was acquired using a commercially available 3i LLS. Briefly, the LLS uses a thin light sheet to achieve single layer excitation for live-cell imaging (4). LLSM has two orthogonal objective lenses: a 0.71 NA, 3.74 mm LWD water-immersion illumination objective and a 1.1 NA, 2 mm LWD water immersion imaging objective, matched to the light sheet thickness for optimal optical sectioning. The lattice creates an evenly illuminated plane of interest, which enables high spatiotemporal resolution of 230 × 230 × 370 nm in xyz. We used a light sheet under a square lattice configuration in dithered mode. Images were acquired with a Hamamatsu ORCA-Flash 4.0 scientific complementary metal-oxide semiconductor (sCMOS) camera. Each plane of imaged volume was exposed for 10 ms with 642 nm laser. The sample was imaged on a piezo stage with the dithered light sheet moving at 276 nm step size in the z-axis. To create an eSRRF ‘frame’, a time-lapse of 100 frames were taken per axial plane.

### Spinning-disc confocal sample preparation & acquisition

To culture **cells on polyacrylamide (PAM)** gel U2OS cells expressing endogenously tagged paxillin-GFP (5) were grown in DMEM/F-12 (Dulbecco’s modified Eagle’s medium/Nutrient Mixture F-12; Life Technologies, 10565–018) supplemented with 10% fetal bovine serum (FCS) (Biowest, S1860). U2OS cells were purchased from DSMZ (Leibniz Institute DSMZ-German Collection of Microorganisms and Cell Cultures, Braunschweig DE, ACC 785). Cells were left to spread on ~9.6 kPa polyacrylamide gel and imaged using a spinning disk confocal microscope. The spinning disk confocal microscope (spinningdisk confocal) used was a Marianas spinning disk imaging system with a Yokogawa CSU-W1 scanning unit on an inverted Zeiss Axio Observer Z1 microscope controlled by SlideBook 6 (Intelligent Imaging Innovations, Inc.). Images were acquired using a Photometrics Evolve, back-illuminated EMCCD camera (512 × 512 pixels). The objective used was a 63× (NA 1.15 water, LD C-Apochromat) objective (Zeiss). 100 frames were used for the eSRRF reconstruction. The parameter sweep option as well as SQUIRREL analyses (resolution-scaled error (RSE) and resolution-scaled Pearson (RSP) values), integrated within eSRRF, were used to define the optimal reconstruction parameters.

The **spheroids** are based on MCF10 DCIS.COM (DCIS.COM) lifeact-RFP cells cultured in a 1:1 mix of DMEM (Sigma-Aldrich) and F12 (Sigma-Aldrich) supplemented with 5% horse serum (16050–122; GIBCO BRL), 20 ng/mL human EGF (E9644; Sigma-Aldrich), 0.5 mg/mL hydrocortisone (H0888–1G; Sigma-Aldrich), 100 ng/mL cholera toxin (C8052–1MG; Sigma-Aldrich), 10 μg/mL insulin (I9278–5 ML; Sigma-Aldrich), and 1% (v/v) penicillin/streptomycin (P0781–100 ML; Sigma-Aldrich). Parental DCIS.COM cells were provided by J.F. Marshall (Barts Cancer Institute, Queen Mary University of London, London, England, UK). To form spheroids, DCIS.com cells expressing lifeact-RFP were seeded as single cells, in standard growth media, at a very low density (3,000 cells per well) on growth factor reduced (GFR) Matrigel-coated glass-bottom dishes (coverslip No. 0; MatTek). After 12 h, the medium was replaced by a normal growth medium supplemented with 2% (vol/vol) GFR Matrigel and 10 μg/ml of FITC-collagen (type I collagen from bovine skin, Merk, Cat Number: C4361). After three days, spheroids were fixed with 4 % PFA for 10 min at room temperature and imaged using a spinning disk confocal microscope. The microscope used as well as the image processing are as described in the previous section.

**Zebrafish** (Danio rerio) housing and experimentation were performed under license MMM/465/712-93 (issued by the Ministry of Agriculture and Forestry, Finland). Transgenic zebrafish embryos expressing mcherryCAAX in the endothelium (genotype Tg(KDR:mcherryCAAX) were cultured at 28.5 °C in E3 medium (5 mM NaCl, 0.17 mM KCl, 0.33 mM CaCl_2_,0.33 mM MgSO_4_). At 2 days post-fertilization, embryos were mounted in 0.7% low-melting-point agarose on glass-bottom dishes. Agarose was overlaid with E3 medium supplemented with 160 mg/l tricaine (Sigma-Aldrich). Imaging was performed at 28.5 °C using a spinning disk confocal microscope. The microscope used as well as the image processing are as described in the previous section with the exception that 150 frames were used for the eSRRF reconstruction.

### Multi-focus microscopy sample preparation & acquisition

HeLa cells were cultured in complete medium (DMEM (11880, Thermo Fisher Scientific) + 1% Glutamax + 1% Penicillin-Streptavidin supplemented with 10% fetal bovine serum (26140079, Thermo Fisher Scientific) and transfected with TOM20 (translocase of outer mitochondrial membrane) fused to HaloTag. TOM20-HaloTag was labeled with Janelia fluor 549 HaloTag ligand (GA1110, Promega) by incubating the dye at 10 nM in DMEM medium for 15 min at 37 °C. MFM imaging was performed in DMEM w/o phenol red medium. The MFM setup used was described in detail in Hajj et al.(6), here excitation was performed with the 555 nm line of a Lumencor Spectra light engine and imaging performed using a Nikon Plan Apo 100x Oil oil immersion objective with NA 1.4. Images of all nine focal planes were captured on an Andor DU-897 EMCCD camera at a rate of 20ms/frame. The focus offset dz was 390 nm between consecutive focal planes. 3D image registration was performed based on multicolor fluorescent beads (TetraSpeck Fluorescent Microspheres Kit; T14792; Invitrogen), immobilized on a coverslip. Images of the beads were recorded while axially displacing the sample with a z-step size of 60 nm. To overlay and align the nine focal planes a calibration table was created based on the bead images with the NanoJ-eSRRF plugin tool “Get spatial registration from MFM data” (NanoJ-eSRRF>Tools>Get spatial registration from MFM data). This spatial registration was applied to the live-cell MFM data during the 3D eSRRF processing (a detailed manual can be found here: https://github.com/HenriquesLab/NanoJ-eSRRF/wiki). To extract the shape of the MFM PSF in the different focal planes (Supplementary Figure 13) the 3D PSF was extracted from the bead reference with the respective NanoJ-eSRRF plugin tool (NanoJ-eSRRF>Tools>Extract 3D PSF from stack).

Deconvolution was performed with the classic maximum likelihood estimation (CMLE) algorithm in the Huygens Professional software by SVI. 3D rendered images were created with Napari (7).

### Estimation of image resolution with FRC with NanoJ-Squirrel(8)

To estimate the image resolution the raw time series image stacks were split into even and odd frames. The independent image sequences were analysed with SRRF, eSRRF or Thunderstorm and based on the resulting pairs of processed images the resolution was estimated by FRC.

## Supplementary Note 1: (e)SRRF and its artifacts

The presence of artifacts in the SRRF reconstruction have been observed previously (1). Here, we highlight an instance where they appear quite clearly (see Supplementary Figure S3). We generated a simulated dataset where the underlying fluorophore arrangement follows a fan configuration. Supplementary Figure S3 shows the reconstructions obtained from SRRF and eSRRF, as well as quantitative measures of image quality based on our SQUIRREL method (2). We observe a clear visual and quantitative improvement of the method. Both resolution-scaled Pearson coefficient (RSP) and resolution-scaled error (RSE) show values associated with improved quality.

## Supplementary Note 2: eSRRF processing and analysis concepts

The eSRRF approach reconsiders the concepts presented in SRRF and uses knowledge of the PSF and the imaging set up to enhance the reconstruction. The method consists in a 2-step process (see Figure 1): a spatial analysis performed on each image of the temporal stack, creating the Radial Gradient Convergence (RGC) stack followed by a temporal analysis on the RGC stack.

### A. Spatial transformation: the Radial Gradient Convergence transform

The SRRF concept exploits the radial symmetry of the signal in the image obtained from a discrete number of emitters in the image. This analysis requires the calculation of image gradients in every single pixel of the image (3). Additionally, the super-resolved image needs to be reconstructed on a finer pixel grid than the acquisition. Therefore, the first step that we take in the spatial reconstruction is to spatially interpolate the raw image by a factor set by the user-set magnification parameter. This is done using Fourier Interpolation via Fast Hartley Transform using JTransforms 3.1, https://github.com/wendykierp/JTransforms. The data is mirror-padded to represent a 2^*n*^x2^*n*^ 2D data block prior to the FHT and the interpolated image is cropped back down to the original image width and height after the interpolation. We have shown that FHT interpolation at an early step of the analysis reduces the occurrence of macro-pixel patterning (Supplementary Figure S1). A set of interpolated frames are sent to the GPU defined automatically by the reconstruction settings. Subsequent calculations are performed on GPU.

The vertical and horizontal image intensity gradients are then calculated from the interpolated images using the Roberts cross method (4).

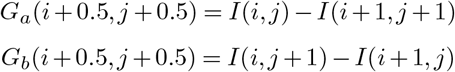

Where *G_a_* and *G_b_* represent the 45 degree angle intensity gradients in the interpolated image. By using Roberts cross, the gradients calculated correspond to those in the corner of each pixel in the original image. The gradients are then rotated by 45 degrees again to be in line with the vertical and horizontal axes of the image using standard 45 degrees rotation matrix calculation, so as to obtain *G_x_* and *G_y_* in each pixel (*i,j*). We found that Roberts cross allows for the estimation of the most local gradients compared to other approaches.

Then the RGC transform is calculated in every pixel of the image. The user-input Radius *R* in pixels represents the FWHM of the expected PSF, thus, the PSF standard deviation can be estimated as follows:

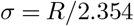

*σ* can be used to calculate the convergence weighting factor *W* as well as the size of the relevant local area over which to calculate the RGC. For a particular pixel of interest the RGC is calculated by summing the weighted gradient convergence (*D_k_*) from all the surrounding pixels in a disk of radius 2*σ*+ 1, called Δ. This radius of calculation allows to speed up the process by only using the relevant local information to the pixel of interest and was determined empirically. The RGC in the pixel (*i*_0_, *j*_0_) is computed as follows:

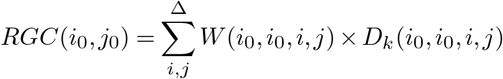

For each pixel in Δ, the distance d to the pixel of interest is computed and used to calculate the weighting factor *W*:

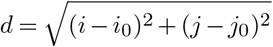

and

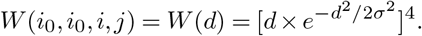

Which is based on the 4-th power of the derivative of a Gaussian pattern. Using the derivative of the Gaussian pattern allows us to weigh more highly the pixels where a strong gradient would be expected if a molecule was present in the pixel of interest. The 4-th power was empirically derived to provide the best local gradient sensitivity.

The dot product of the gradient vector in the adjacent pixel and the distance vector between the adjacent and pixel of interest is computed to know the orientation of the gradient vector with respect to the pixel of interest. If the gradient vector is pointing towards the pixel of interest, then the cross product of the distance vector and the gradient vectors is computed to calculate the distance of the tangent, similar to what was previously done with SRRF. This is used to calculate the gradient convergence for a particular location pair:

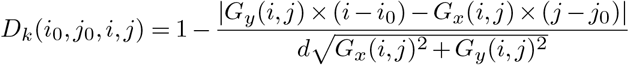

This essentially computes the smallest distance between the gradient vector in (*i, j*) and the point of interest (*i*_0_, *j*_0_), normalized by the distance between these 2 points. *D_k_* then becomes 1 if the gradient points exactly at (*i*_0_, *j*_0_), and decreases as the vector points further and further away.

NB: the RGC grid and the gradient grid are effectively on the same sized grid but the gradients are computed on the corner of the pixels whereas the RGC (and interpolated intensities) are computed on the centre of the pixels.

### B. Temporal transformation: temporal cross-correlation analysis

eSRRF provides three different temporal transformation strategies that each perform best in different emitter density and fluctuation regimes (see Supplementary Figure S5). A temporal average projection (AVG) of the RGC map for each pixel (*i*_0_, *j*_0_) provides robust results with low sensitivity to noise, over a wide range of emitter densities.

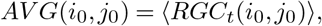

〈…〉 indicates the average over time. However, an additional resolution improvement can only be achieved by higher-order temporal correlations. The temporal variance projection (VAR) which corresponds to cross-cumulant of 1st order without temporal offset and the 2nd order temporal auto-cumulant (TAC2) which corresponds to the cross-cumulant of 1st order with offset of 1 frame (5) provide an additional resolution gain with significant improvements in fidelity and contrast by analyzing the temporal fluctuations:

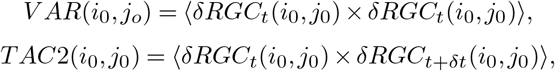

with *δRGC_t_*(*i*_0_, *j*_0_) = *RGC_t_*(*i*_0_, *j*_0_) – 〈*RGC_t_*(*i*_0_, *j*_0_)〉.

From an implementation point of view, the GPU calculates both the temporal average of the RGC maps and the temporal average of the square of the RGC maps. The final VAR is then computed on CPU as follows:

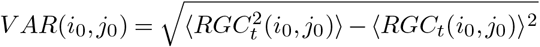

A similar approach is taken for TAC2.

## Supplementary Tables

**Table S1.** Comparison of fluctuation-based super-resolution microscopy methods.

| Method | Reference | Basic principles | Implementation | GPU acceleration | Image fidelity validated | 3D | In depth |
| --- | --- | --- | --- | --- | --- | --- | --- |
| eSRRF | this publication | Radiality and temporal cross-correlation | Fiji | Yes | Yes | Yes | Yes |
| SRRF | Gustafsson *et al.*, 2016(6) | Radiality and temporal cross-correlation | Fiji | Yes | No | No | Yes |
| MUSICAL | Agarwal *et al.*, 2016(7) | Multiple signal classification | Fiji | Yes | No | No | Yes |
| SOFI | Dertinger *et al.*, 2009(8) | Temporal cross-correlation | MATLAB | No | No | Yes | Yes |
| SACD | Zhao *et al.*, 2020(9) | Deconvolution, temporal cross-correlation | MATLAB | No | No | No | ? |

**Table S2.** eSRRF parameters.

| Parameter | Description |
| --- | --- |
| Magnification $M$ | Define how the camera pixels are split into sub-pixels for the RGC estimation. |
| Radius $R$ | Define the receptive field size that is used to calculate the RGC, the size should represent the FWHM of the expected PSF. |
| Sensitivity $S$ | Define the sensitivity factor to fine-tune the PSF sharpening power applied by the RGC. |
| Number of frames for eSRRF | Define the size of the frame window for the temporal analysis. |
| Vibration correction | Activate vibration correction based on cross correlation. |
| Temporal analysis method | Select AVG, VAR and/or TAC2 as a temporal analysis method. |
| Rolling analysis | Activate rolling analysis and define frame gap size. |
| 3D eSRRF | Activate 3D eSRRF analysis and define offset between axial planes in nm. |

**Table S3.**
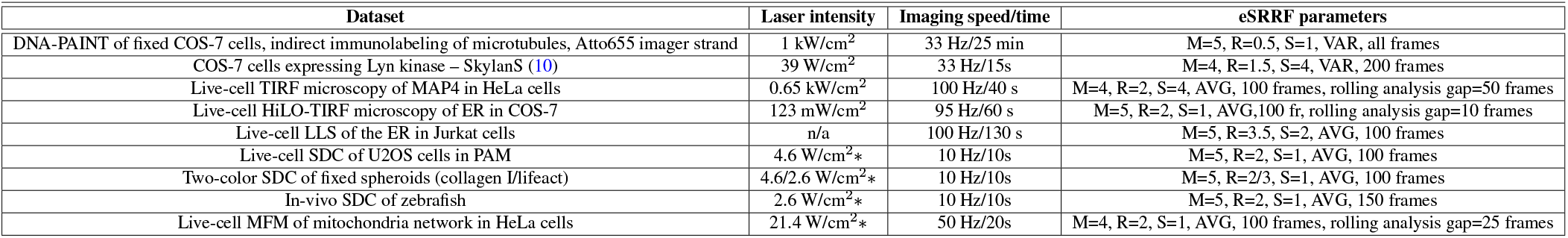
Laser intensities, imaging speed, total imaging time and eSRRF parameters for the data sets included in this paper. *Intensity at the coverslip surface.

## Supplementary Figures

**Fig. S1.**
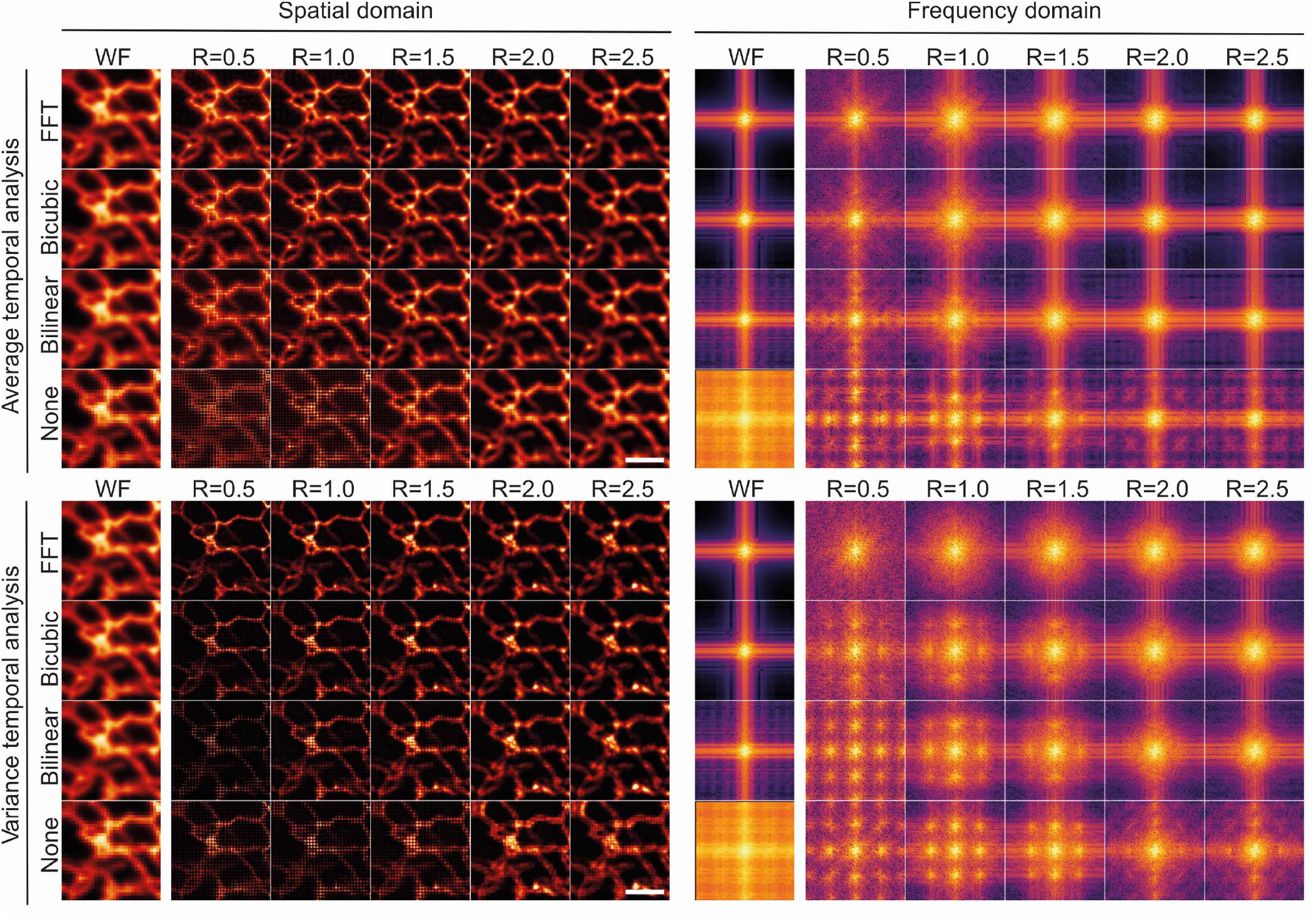
Comparison of interpolation methods in the reconstruction of SRRF images. The presence of macro-pixel artifacts is apparent in both the spatial (left column), and frequency domain (right column) and for AVG (upper row) and VAR temporal analysis reconstruction (lower row) for all interpolation methods apart from the FFT-based interpolation. Data shown corresponds to live COS-7 cells expressing PrSS-mEmerald-KDEL acquired using HiLO-TIRF. Scale bars 2 μm.

**Fig. S2.**
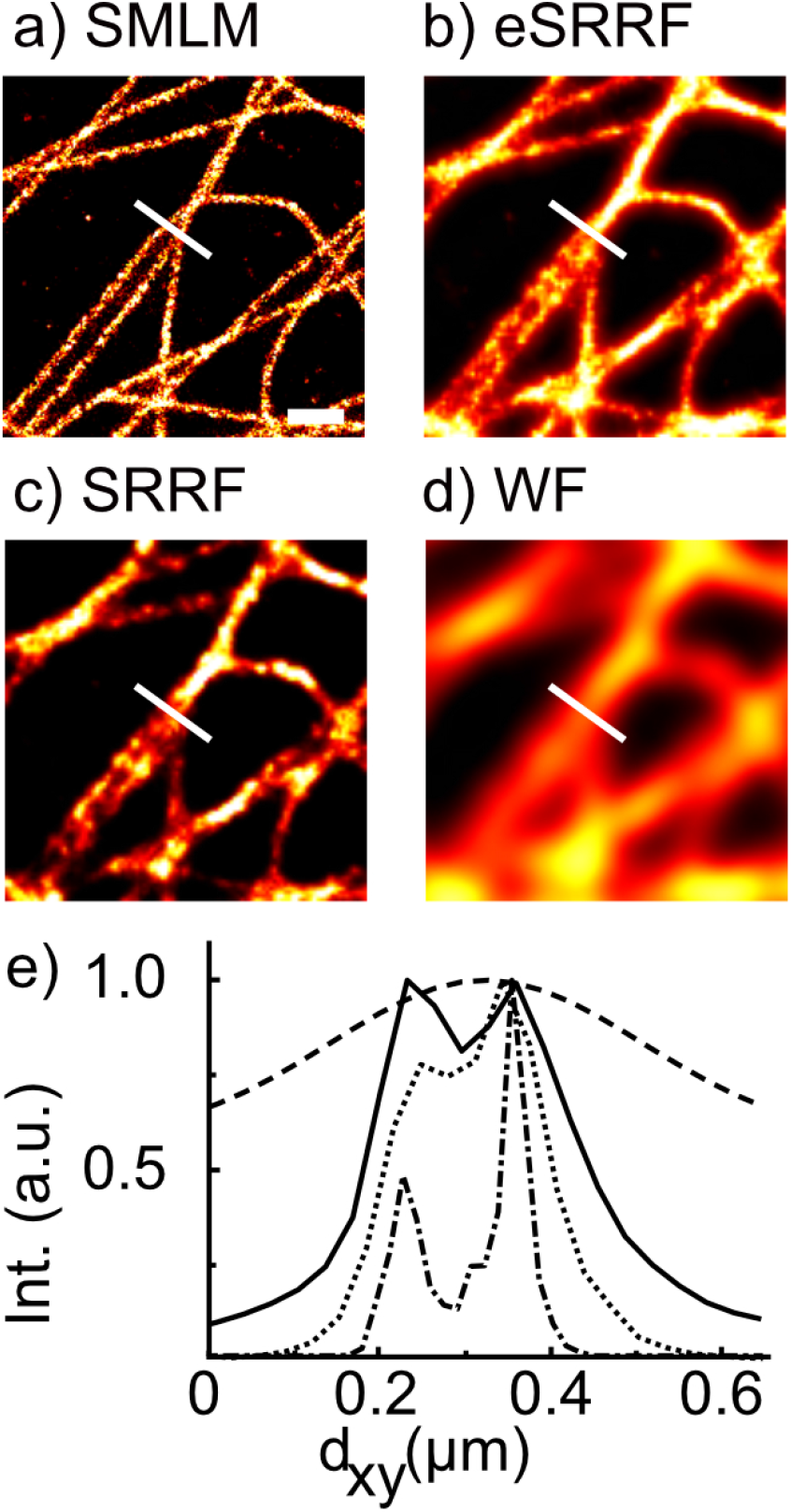
Resolution improvement of eSRRF vs. SRRF. Image sections of the data set presented in Figure 1 after a) SMLM image reconstruction, b) eSRRF processing, c) SRRF processing and as WF data. The white line indicates the position of the line profiles. e) Intensity profiles allow to distinguish two filaments in the SMLM reconstruction (dash-dotted line) which can also be resolved with eSRRF (solid line) but the presence of a second filament is unclear in the case of SRRF processing (dotted line) and for the WF data (dashed line). Scale bar: 500 nm.

**Fig. S3.**
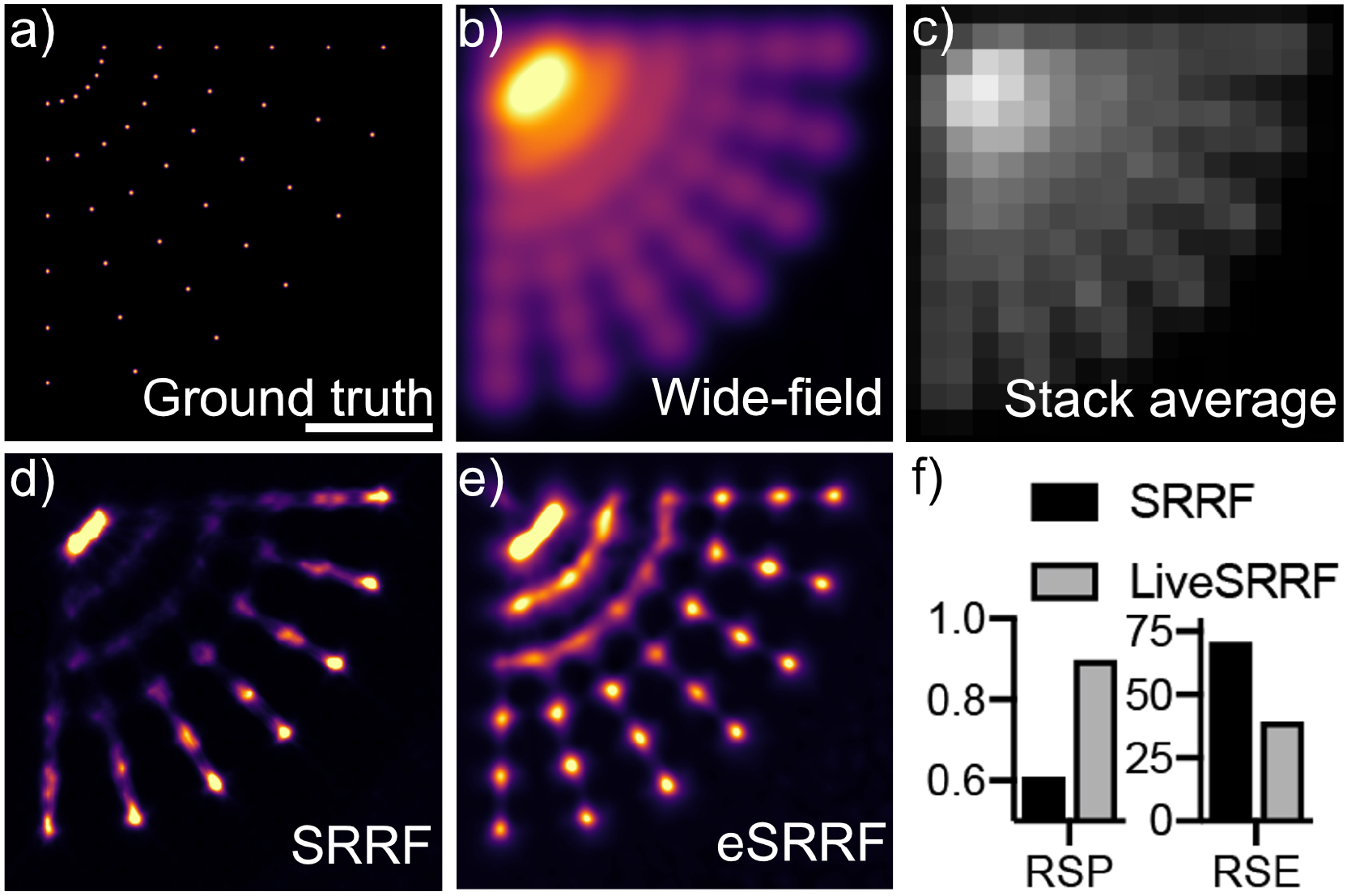
Performance improvement of eSRRF on simulated data. a) Simulated ground truth indicating the positions of individual molecules in the image. b) Interpolated wide-field image. c) Average of all simulated raw frames. d) SRRF image. e) eSRRF image. f) Quantitative comparisons of SRRF and eSRRF based on RSP and Resolution Scaled Error (RSE) obtained from SQUIRREL. Artifacts like the linearity loss and and over sharpening as they are observed in d) are significantly reduced with e) eSRRF processing, Scale bar: 500 nm.

**Fig. S4.**
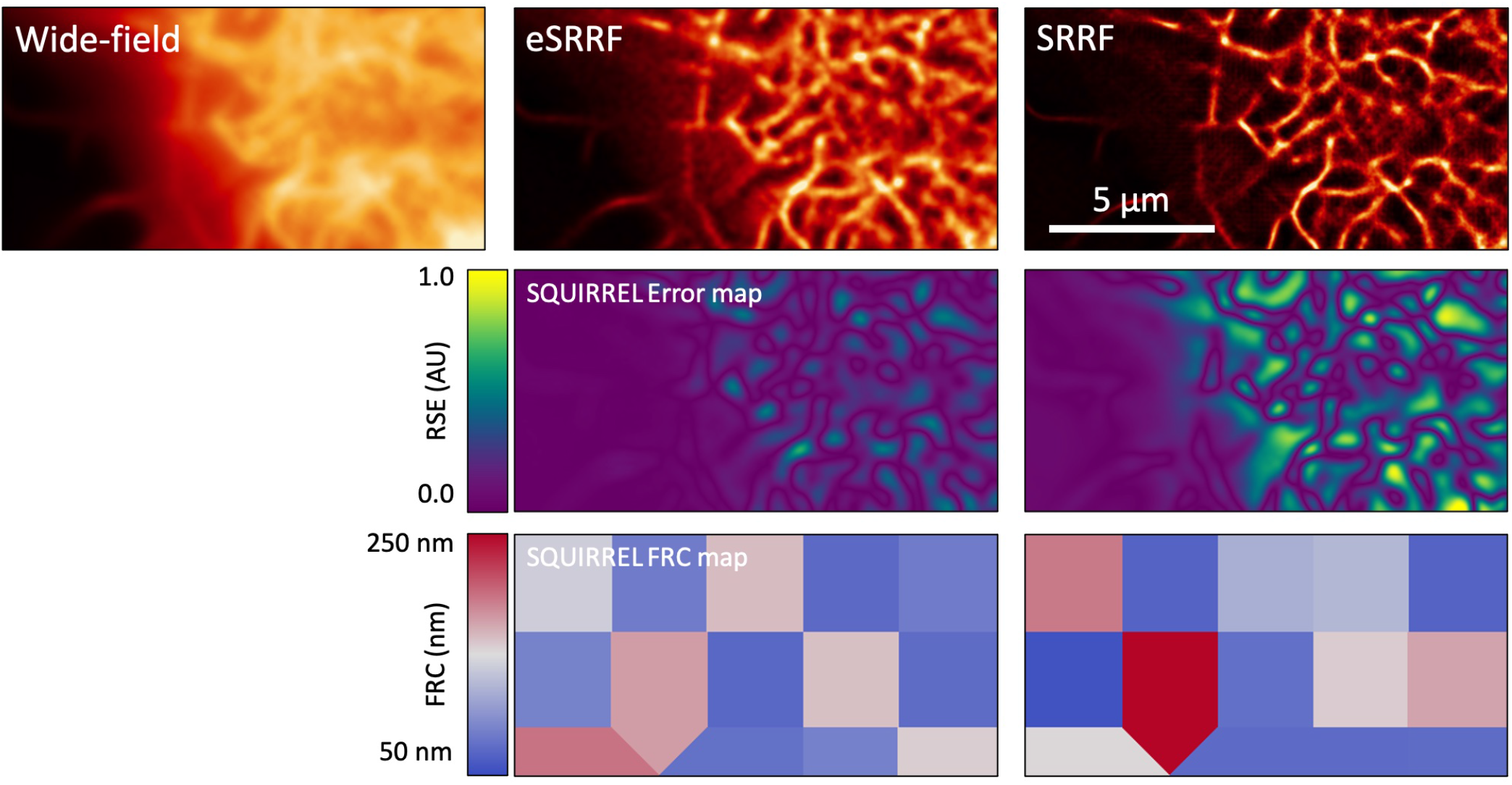
SQUIRREL comparison of SRRF and eSRRF. eSRRF of the actin network (GFP-UtrCH expression, 100 frames at 33 frames/s) in live COS-7 cells shows an improved fidelity with a retained FRC resolution range. The dataset was published before in Culley et al.(11)

**Fig. S5.**
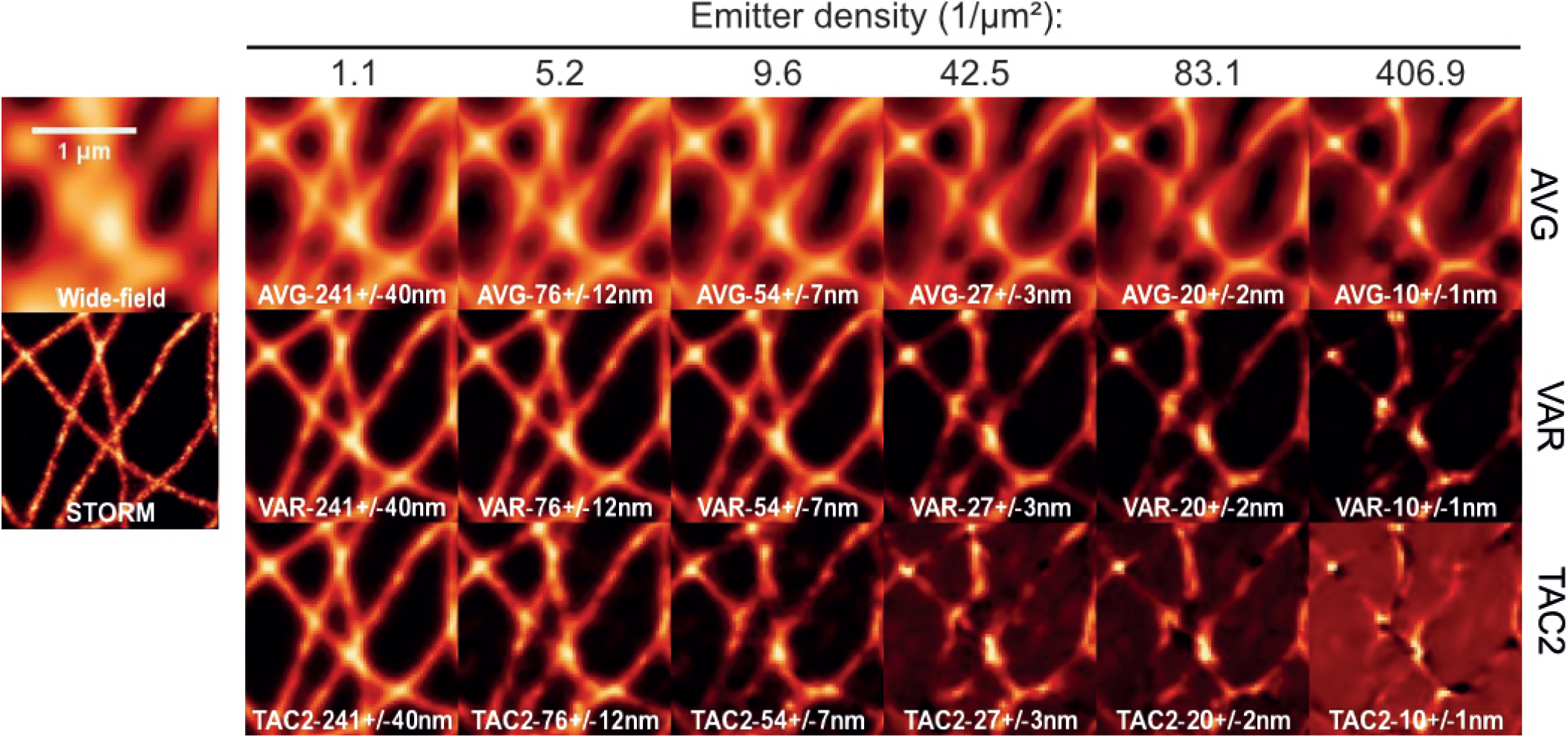
Temporal analyses of eSRRF as a function of the density of emitters in the raw data images. The reconstructed images are shown in increasing emitter density from left to right. The corresponding average nearest-neighbor distance is stated in each panel. The wide-field and STORM equivalents are shown on the left for comparison. Scale bar 1 μm.

**Fig. S6.**
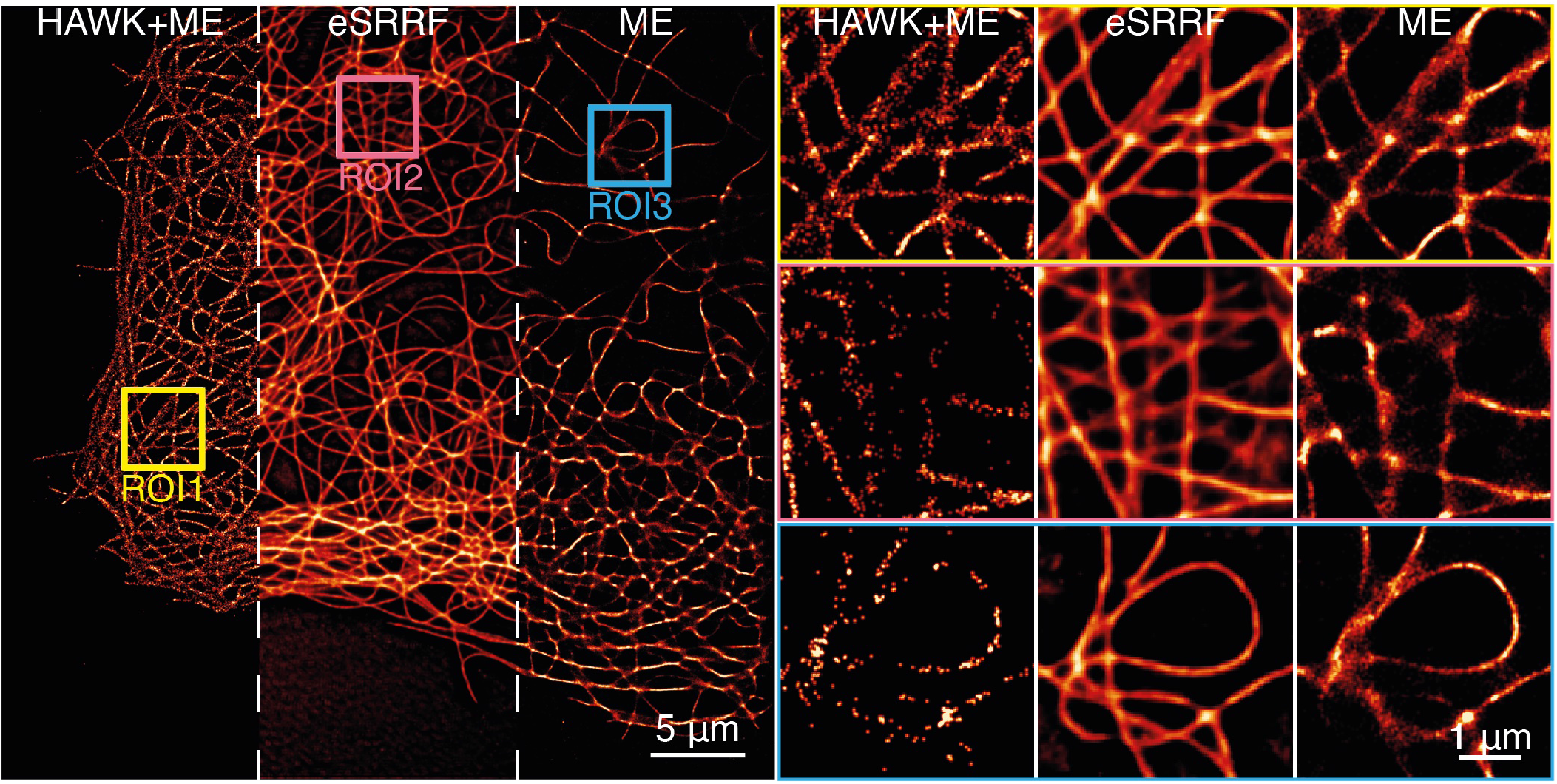
Comparison of eSRRF, HAWK+ME and ME. The Maximum Likelihood Estimation multi-emitter fitting (ME) was performed using ThunderSTORM(12). Data shown corresponds to a DNA-PAINT acquisition of immunolabeled microtubules in fixed COS7 cells under TIRF illumination.

**Fig. S7.**
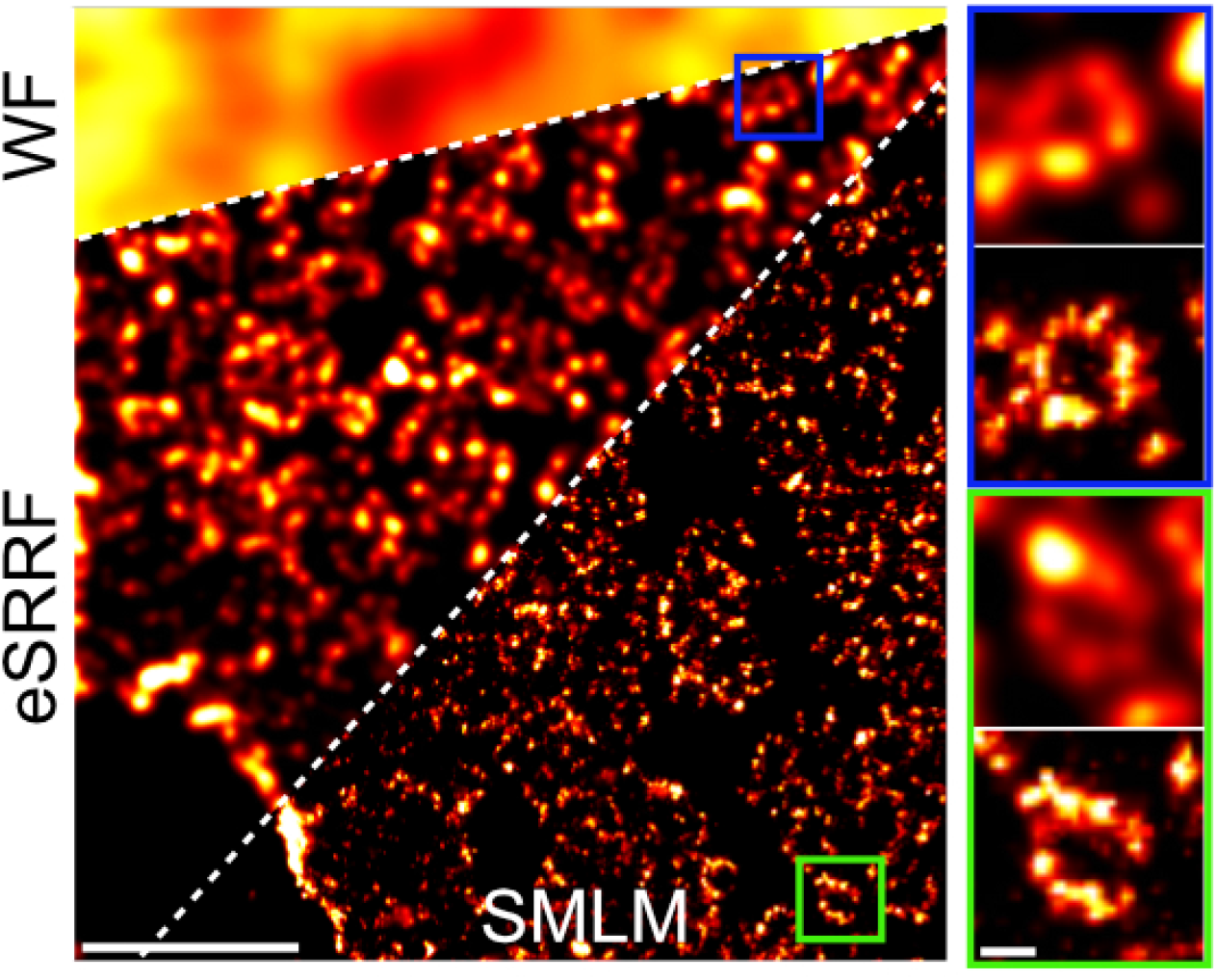
eSRRF allows a fast preview of SMLM dataset reconstruction. Widefield imaging (top left) of the nuclear pore complex. Fast eSRRF reconstruction preview (first 2000 frames shown) reveals the open ring structure (left panel middle area and insets top panels). While fast eSRRF can resolve the central pore which has a diameter of about 140 nm, the full 8-element ring with only 40 nm gaps is only resolved by single-molecule localization analysis of the full 20 000 frames image stack (left panel bottom area and insets lower panels. The FRC resolution is 44.4±2.5 nm, and 35.1±6.3 for eSRRF and SMLM, respectively.,) (Dataset from Heil et al. (13)), left panel: scale bar 1 μm, insets on the right: scale bar 100 nm.

**Fig. S8.**
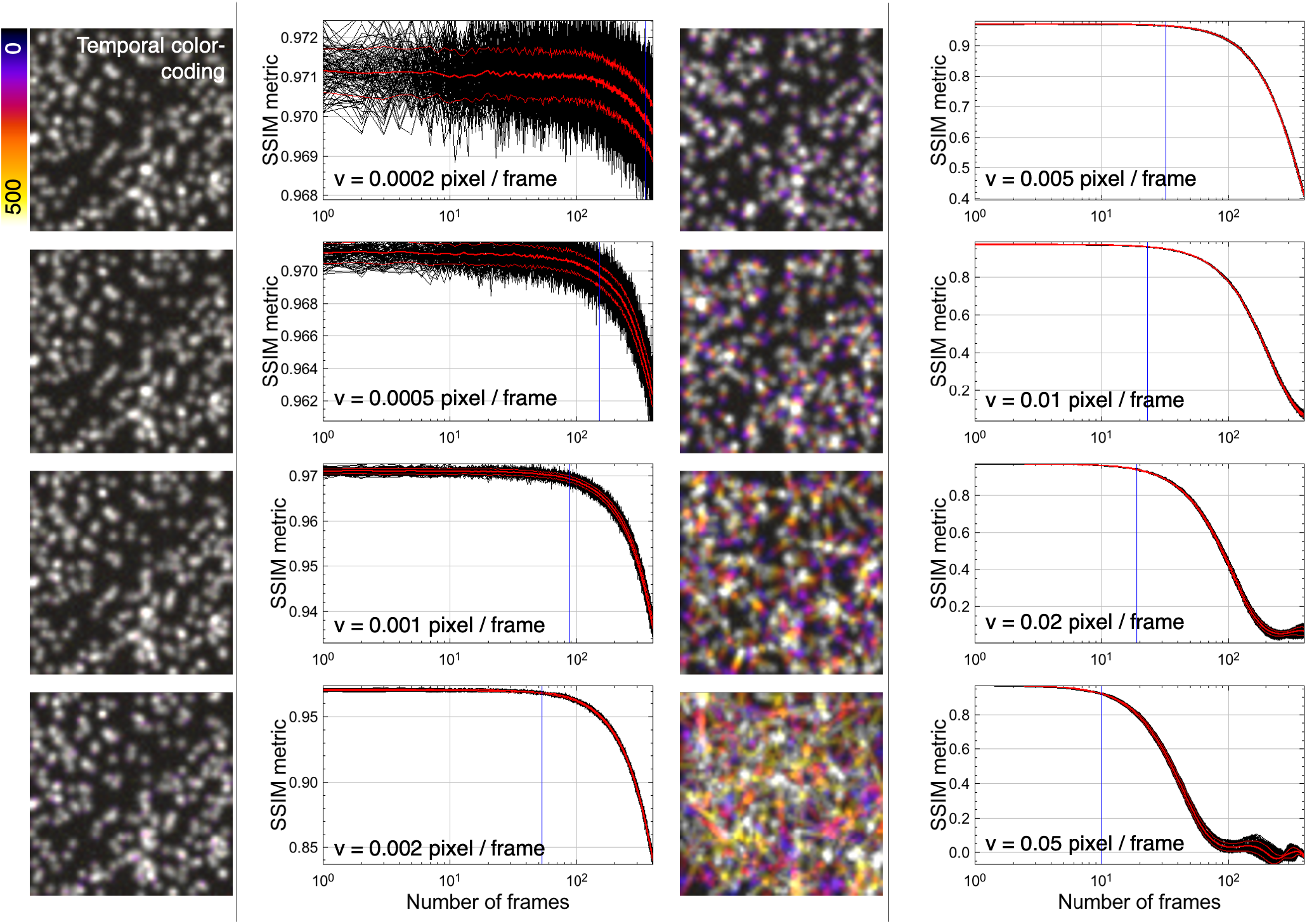
tSSIM analysis of simulated moving particles. Color coded projections of simulated image stacks displaying particles diffusing with various speeds v and the resulting SSIM metric progression over time. The tSSIM metric shows sensitivity as a function of particle displacement per frame. This can be used to estimate the size of the optimal time window for eSRRF processing to avoid movement artifacts.

**Fig. S9.**
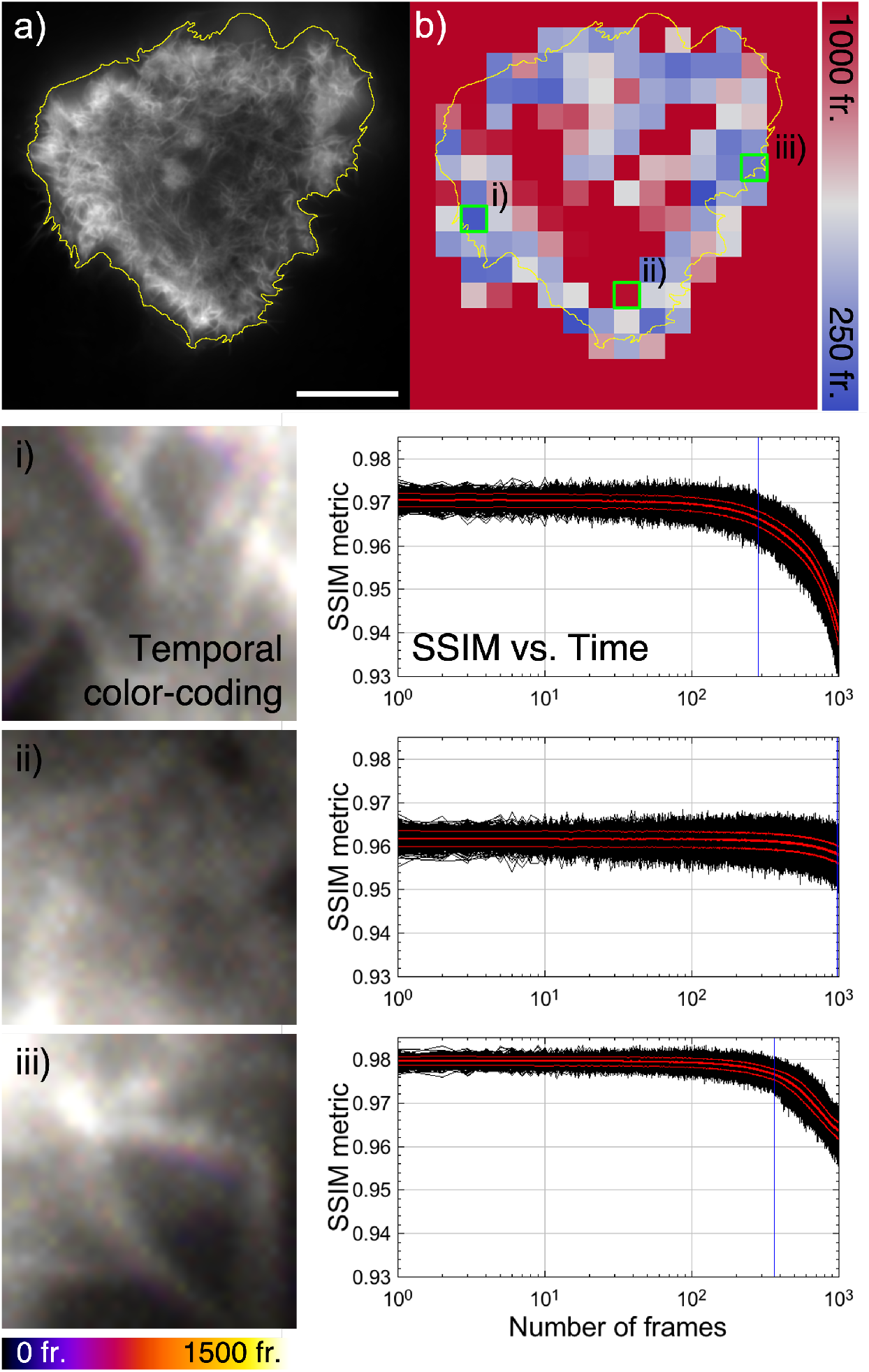
tSSIM analysis performed over small image patches detects local dynamics of actin rearrangement in COS-7 cells. a) The actin network in live COS-7 cells expressing the marker GFP-UtrCH was acquired at 33 fps. b) The tSSIM metric estimates the time range of motion within individual subsections of a). The different subsections highlighted in green display the color-coded projection of regions with i) fast, ii) slow, and iii) moderate speeds as is also reflected by the corresponding progression of the SSIM metric over time. Scale bar 20 μm.

**Fig. S10.**
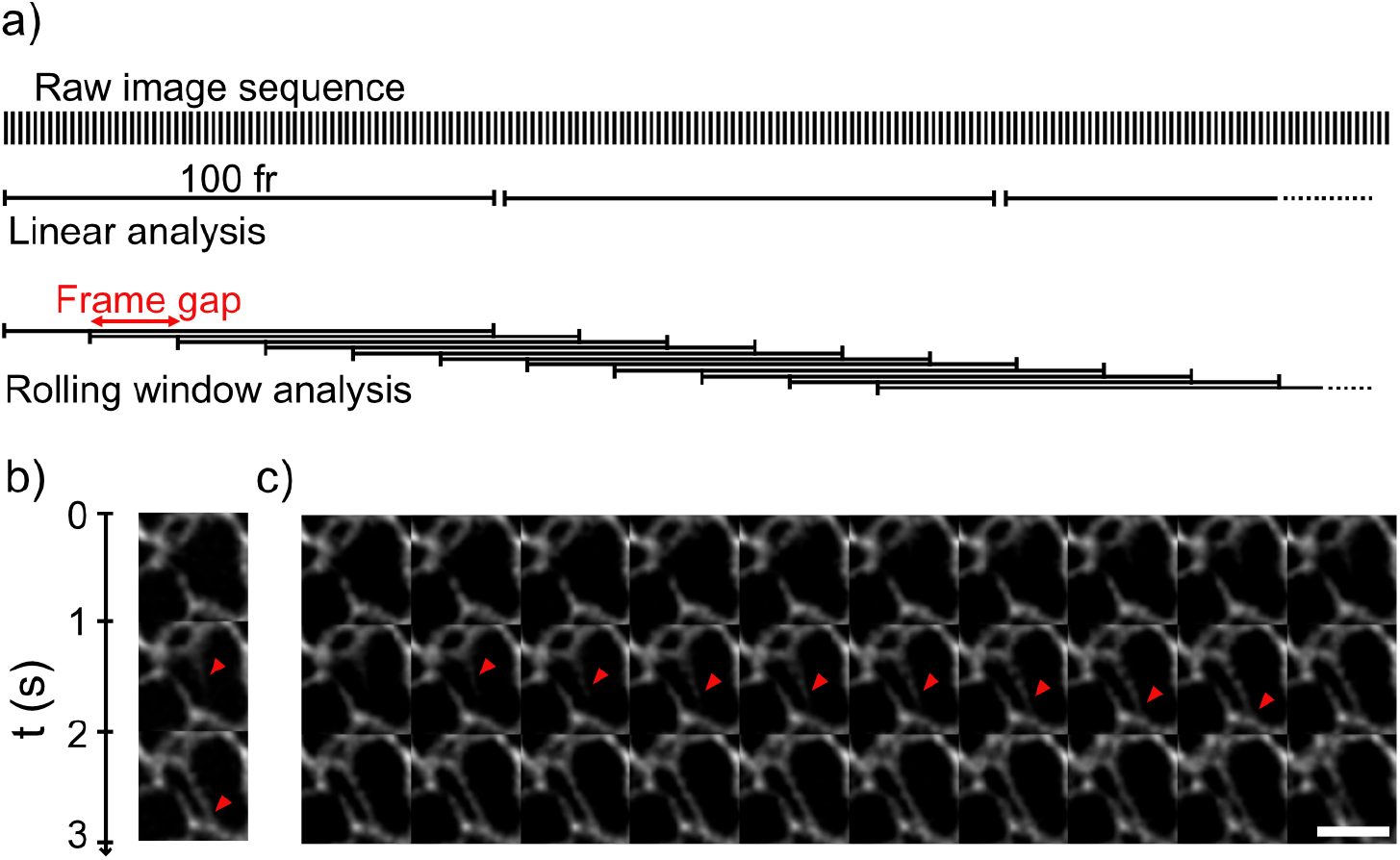
Increasing temporal sampling by rolling window analysis. a) Super-resolved temporal image stack is reconstructed by analyzing consecutive frame windows (linear analysis). A rolling window analysis with a frame gap of less than the window size allows to increase the temporal sampling rate and can translate into a higher temporal resolution without sacrificing spatial resolution. b) This allows to visualize dynamic rearrangement of the endoplasmic reticulum (ER, red arrow) acquired by live-cell HiLO-TIRF of COS-7 cells expressing PrSS-mEmerald-KDEL at a frame rate of 1 Hz with a 100 frame-window linear eSRRF analysis. c) The rolling window analysis with a frame gap of 10 frame allows to increase the temporal sampling to 10 Hz, thus, revealing the substeps of the ER tubule formation (red arrow), image section width 3 μm, FRC resolution: eSRRF: 143±56 nm, Scale bar 2 μm.

**Fig. S11.**
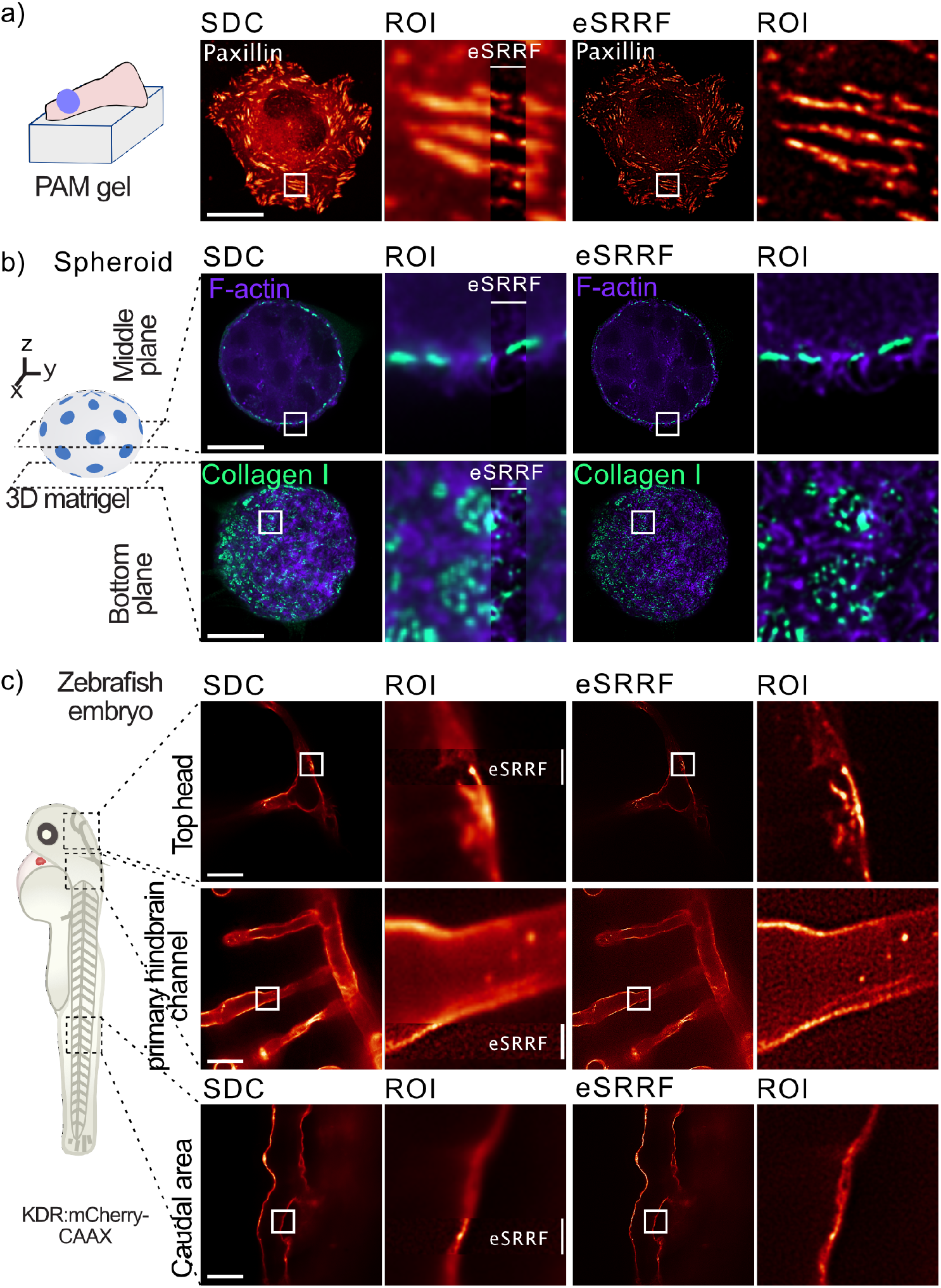
eSRRF enhances spinning-disk confocal imaging deep inside fixed and live cells, spheroid and organisms. a) U2OS cells expressing endogenously tagged paxillin were plated on 9.6 kPa polyacrylamide (PAM) gels and were imaged live using a spinning-disk confocal (SDC). b) DCIS.com lifeact-RFP cells forming a spheroid in 3D matrigel in the presence of fluorescently labeled collagen I. Samples were fixed and imaged using a spinning-disk confocal microscope and processed using eSRRF. Representative fields of view highlighting the spheroids’ middle and bottom planes are displayed. c) Zebrafish embryos expressing mcherryCAAX in the endothelium were imaged live using a spinning-disk confocal. Vessels located at different parts of the embryo were imaged. For all panels, the eSRRF reconstructed images and the original spinning disk images are displayed. Scale bars 25 μm.

**Fig. S12.**
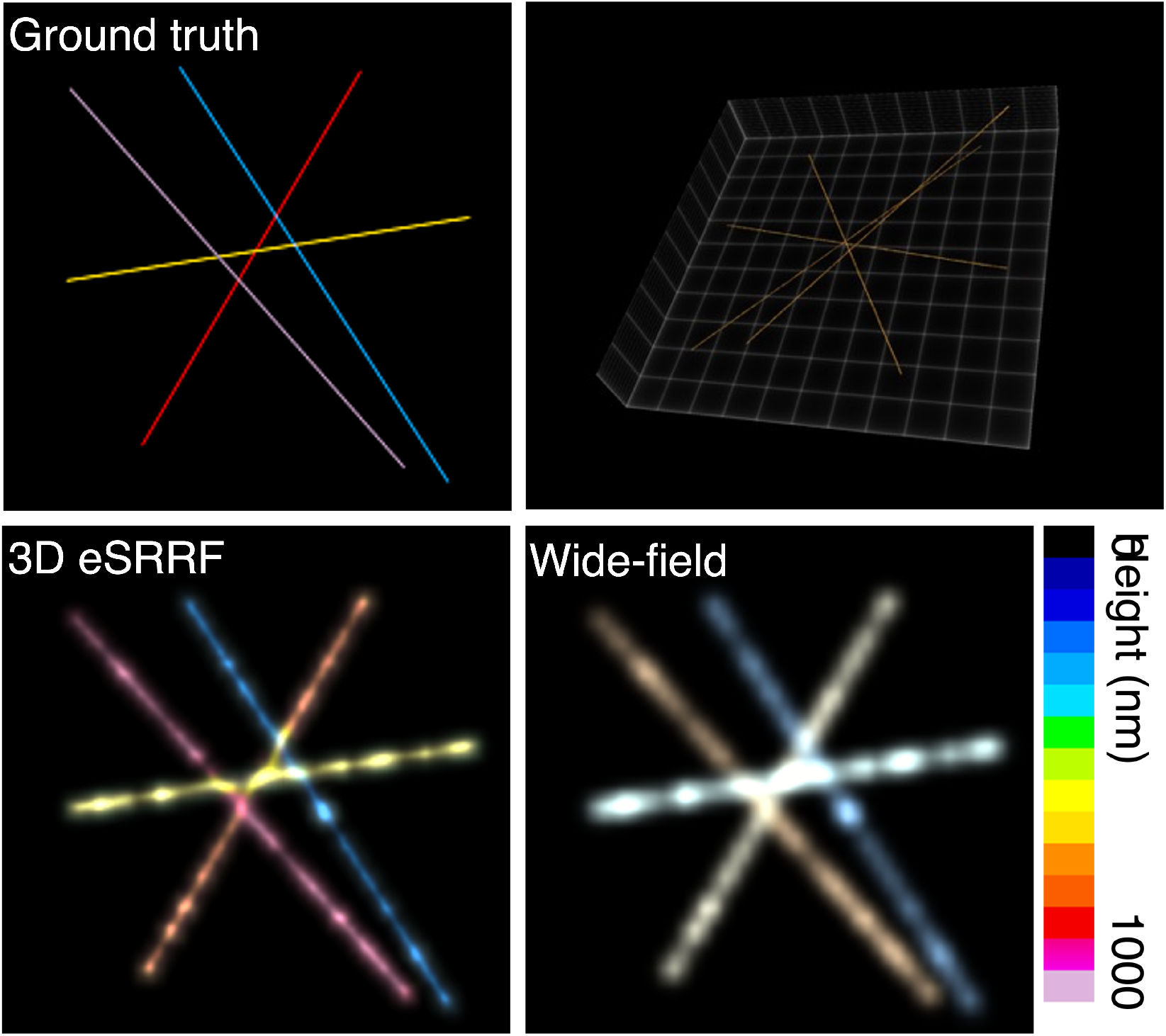
3D eSRRF on a simulated dataset. The height of the filament is color coded as indicated by the false color scale.

**Fig. S13.**
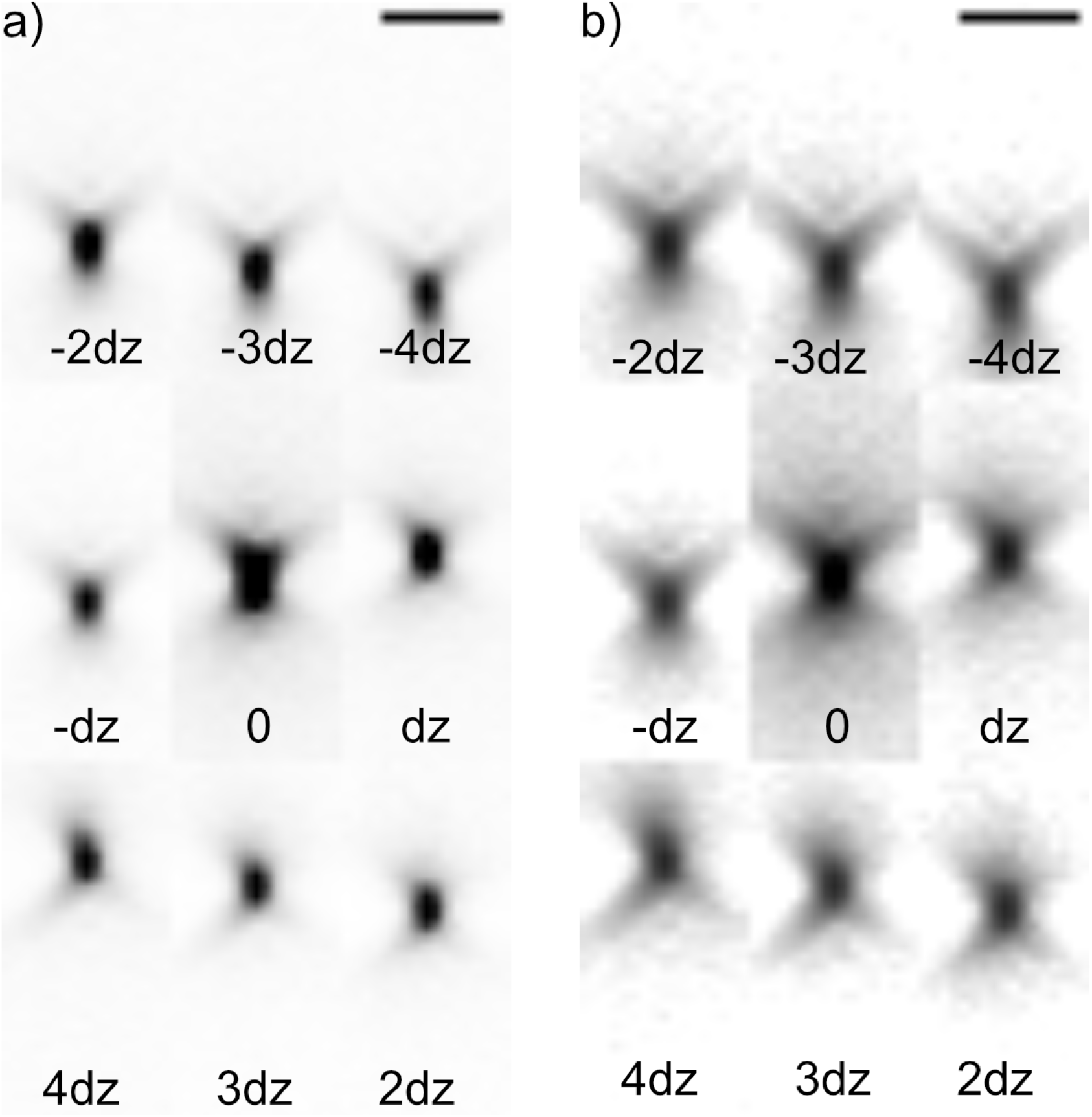
3D PSF in the nine focus planes of the MFM displays only minor aberrations and good radial symmetry. x-z view of the PSF mapped with the bead calibration dataset displayed in a) linear and b) logarithmic brightness scale (FWHMX=431±19 nm, FWHMz=704±45 nm). The focus offset dz between each focal plane is 390 nm. Scale bars 2 μm.

**Fig. S14.**
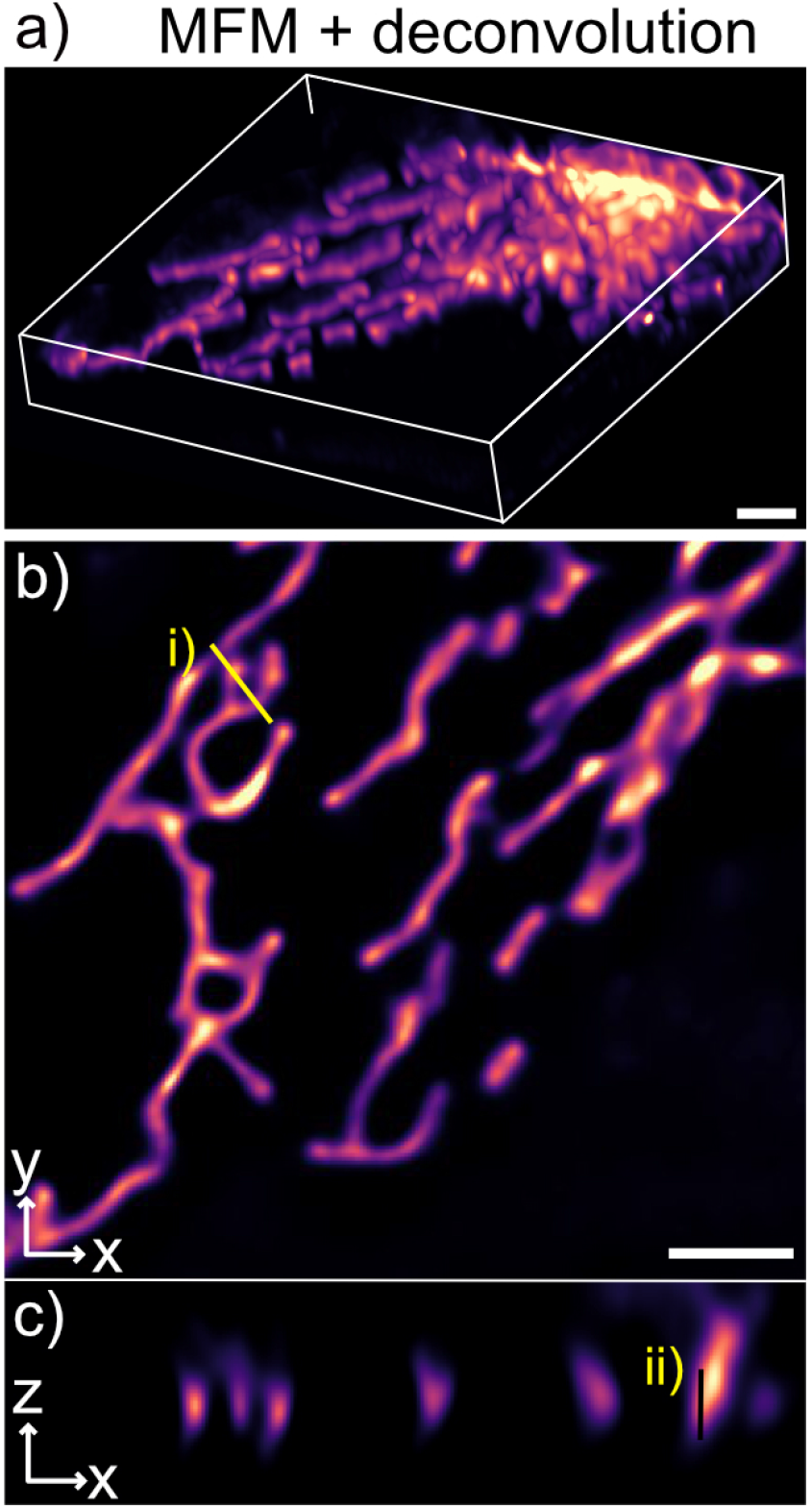
Deconvolved 3D image stack of U2OS cells expressing TOM20-Halo, loaded with JF549 acquired with MFM. a) 3D rendering, b) single z-slice and c) single cropped y-slice. i) and ii) are the line profiles in x,y and z-plane displayed in Figure 4. Scale bars 2 μm.

## Supplementary Movies

**Supplementary Movie M1: Automated reconstruction parameter search tool implemented in eSRRF.** 200 frames of the live-cell TIRF imaging dataset of COS-7 cells expressing Lyn kinase – SkylanS were analysed with eSRRF covering the Radius R and Sensitivity S parameter space defined by *R_start_* =1, step size=0.5, number of steps=7 and *S_start_* =1, step size=1, number of steps=8. The eSRRF reconstruction for each parameter combination is presented on the left, while the corresponding image resolution and fidelity is marked with a yellow square in the respective FRC and RMSE maps. At a low R values pixel artifacts are evident, while at higher R values and low S values no high resolution is achieved. If both, R and S values, are high the reconstruction displays a high degree of nonlinearity. The compromise between resolution and fidelity is represented in the QnR map which displays a maximum at the parameter combination R=2 and S=4 (marked in red).

**Supplementary Movie M2: Live-cell HiLO-TIRF of COS-7 cells expressing PrSS-mEmerald-KDEL marking the ER lumen.** WF and eSRRF reconstruction of COS-7 cells expressing a luminal ER marker allows live-cell super-resolution imaging (FRC resolution WF/eSRRF: 254±11/143±56 nm) at a sampling rate of 1 Hz. Rolling window analysis allows to speed up temporal sampling to 10 Hz.

**Supplementary Movie M3: Lattice-light sheet imaging of ER in live Jurkat T-cells enhanced by eSRRF.** Slice-by-slice processing of the data set allows the reconstruction of a volumetric view (79 x 55 x 35 μm^3^) of the ER network in live Jurkat T-cells at a rate of 7.6 mHz.

**Supplementary Movie M4: Live-cell 3D eSRRF of mitochondria dynamics with MFM.** Live-cell volumetric imaging of U2OS cells expressing TOM20-Halo, loaded with JF549 with MFM of a 20 x 20 x 3.6 μm3 at a volume frequency of 1Hz, the grid size is 2 μm.

**Supplementary Movie M5: Observing mitochondria dynamics with live-cell 3D eSRRF with MFM.** Single z slice image over the whole MFM acquisition time of U2OS cells expressing TOM20-Halo loaded with JF549, before (left) and after eSRRF processing (right). Scale bar 2 μm.

## Notes

https://github.com/HenriquesLab//NanoJ-eSRRF

https://doi.org/10.5281/zenodo.6466472

### Competing Interest Statement

The authors have declared no competing interest.

### Summary of Updates

Author list update

